# Low oscillatory shear stress regulates Weibel-Palade body size and vWF release

**DOI:** 10.1101/2023.05.26.542527

**Authors:** Ashley Money, Harriet J Todd, Hanjoong Jo, David J Beech, Lynn McKeown

## Abstract

Blood flow regulates vascular function by generating wall shear stress which impacts endothelial cell (EC) physiology. Whilst high laminar shear stress (HSS) maintains the functional integrity of the vasculature, low oscillatory shear stress (LOSS) evokes secretion of pro-thrombotic and pro-inflammatory components from ECs, thus promoting the development of cardiovascular disease (CVD). Pro-thrombotic and pro-inflammatory cargo are readily stored in endothelial-specific organelles termed Weibel-Palade bodies (WPBs). WPBs form purely due to the multimerization of the pro-thrombotic glycoprotein von Willebrand factor (vWF). Here we investigated if aberrant shear stress, induced by non-uniform oscillatory flow, modulates the biogenesis of WPBs, and the subsequent release of vWF. Ultimately, we demonstrate, using an *in vitro* model, that LOSS has no effect on the level of vWF expression, however, LOSS promotes endothelial thrombotic potential by increasing WPB size, which impacts the length of the vWF strings that are released upon stimulation. Thus, wall shear stress can confer functional plasticity to WPBs by manipulating their biogenesis. Further understanding of the underlying mechanisms could aid generation of novel therapeutics in CVD.

## Introduction

Weibel-Palade bodies (WPBs) are endothelial cell (EC)-specific organelles of 1-5 µm in length generated wholly by the production and multimerization of von Willebrand factor (vWF) ^1 2 3^. Often termed ‘vascular first aid kits’, WPBs store and release ingredients that enable healing in response to vascular injury. The stimulated release of vWF filaments from ECs into the vessel lumen is necessary for attracting platelets to the site of damage in order to form a plug that blocks excessive bleeding. Simultaneously, WPB mediated surface exposure of P-selection attracts leukocytes for removal of bacteria ^4^, and the release of angiopoietin-2 (Angpt2) promotes endothelial migration and wound closure ^5^. Storage of ready-made cargo primed for rapid release is a necessary but risky strategy for the endothelium; indeed, increased plasma levels of vWF contributes to the development of atherosclerotic plaques and thus cardiovascular disease ^6 7 8^. For example, dysregulated secretion of vWF promotes atherosclerotic plaque formation by inappropriate aggregation of platelets ^9^, vWF deficient animal models show decreased plaque formation and less coronary thrombi ^10 11^ and a large study in the Netherlands demonstrated that patients with diabetes that are deficient in vWF have less cardiovascular events ^12^.

Fluid shear stress, a frictional force generated by blood flow, plays a role in the physiological functions of ECs ^13^. Unidirectional high laminar shear stress as sensed by ECs in straight vessels is atheroprotective and inhibits inflammatory gene expression via KLF2 (Kruppel-like factor2), a laminar shear stress-induced zinc-finger transcription factor. In contrast, areas of low or disturbed shear stress under oscillatory flow, such as vessel branch points, bifurcations and curvatures, show altered patterns of inflammatory gene expression including increased expression and activation of NFκB (a transcription factor responsible for expression of pro-inflammatory genes under pathogenic flow) and promote the development of atherosclerotic plaques ^14^ Moreover, shear stress may regulate release of WPB cargo. Galbusera et al. demonstrated that sudden exposure of static ECs to high shear evokes exocytosis of vWF ^15^, whilst Sun et al ^16^ revealed that the vWF secretion evoked by short term exposure to high shear is not maintained at longer time periods unless shear stress is lowered. In addition, mimicking high shear laminar flow by heterologous expression of KLF2 attenuates thrombin induced vWF release from WPBs in bone-derived progenitor ECs ^17^. These data are consistent with vWF being localised only to areas subjected to low shear or non-uniform flow in vivo ^10,18^.

The mechanisms underlying the impact of shear stress on WPBs are not well understood. Whilst release of specific cargo can be regulated at the cell surface ^19 20 21 22^, it is also becoming increasingly clear that ECs can generate subpopulations of WPBs, carrying distinct cargo, in response to environmental cues ^23 24 25^. Thereby, we reasoned that shear stress is a modulator of WPB biogenesis. To test this idea we observed WPB biogenesis and vWF recruitment in Human Umbilical Vein Endothelial Cells (HUVECs: used as a robust model of WPB function) and Human Aortic Endothelial Cells (HAECs: used as a flow-responsive and physiologically relevant model of arterial ECs). ECs were cultured under physiological high laminar shear stress (HSS) and compared to static or low oscillatory shear stress (LOSS) conditions. We hypothesize that HSS and non-uniform LOSS has differential effects on the biogenesis of WPBs thereby affecting cargo recruitment and vWF release.

## Methods

### Cell Culture

HUVECs (PromoCell) were maintained in Endothelial Cell Basal Medium (EBM-2) supplemented with Endothelial Growth Medium-2 Supplement Pack (PromoCell). Pooled HAECs (PromoCell) were maintained in Endothelial Cell Growth Medium MV2 supplemented with Growth Medium MV2 Supplement Pack (PromoCell). All cells were maintained at 37°C and 5% CO_2_ in a humidified incubator. Cells were grown to confluency before passage and used between passages 1-5.

### Ibidi Pump System

ECs were plated on Ibidi µ-slide I^0.4^ Leur (IbiTreat) or Ibidi y-shaped slides (IbiTreat) at a concentration of 7.5 × 10^5^ cells/ml and incubated at 37°C and 5% CO_2_ until confluent (18-24 h). To subject the cells to flow, each Ibidi slide was connected to a fluidic unit, with reservoirs containing 15 ml EGM-2. Based on the parameters input into the Ibidi software, the media was flowed over the cells at different rates of shear stress. Flow was applied in either a continuous unidirectional manner (HSS) or alternating direction every second due to electrically controlled valves within the fluidic unit (LOSS) on the Ibidi µ-slides. In this instance, HUVECs were exposed to either high unidirectional shear stress (HSS; 10 dyn/cm^2^), low oscillatory shear stress (LOSS; 2 dyn/cm^2^, oscillating at 1 Hz) or static conditions for 48 hours concomitantly with daily media change on the static slide.

### RT-qPCR

Total RNA was extracted from fresh cells using high pure RNA isolation kit (Roche) as per manufacturer’s instructions. Complimentary DNA (cDNA) was synthesised using a High Capacity RNA-to-cDNA RT kit (Applied Biosystems). RT-qPCR was carried out using SYBR Green I (Bio-Rad) and a LightCycler (Roche). Primers are provided in Table 1.

**Table 1:**
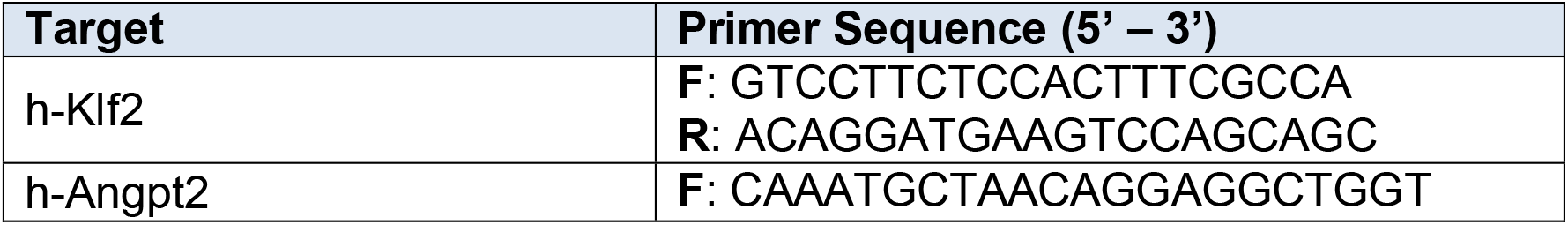
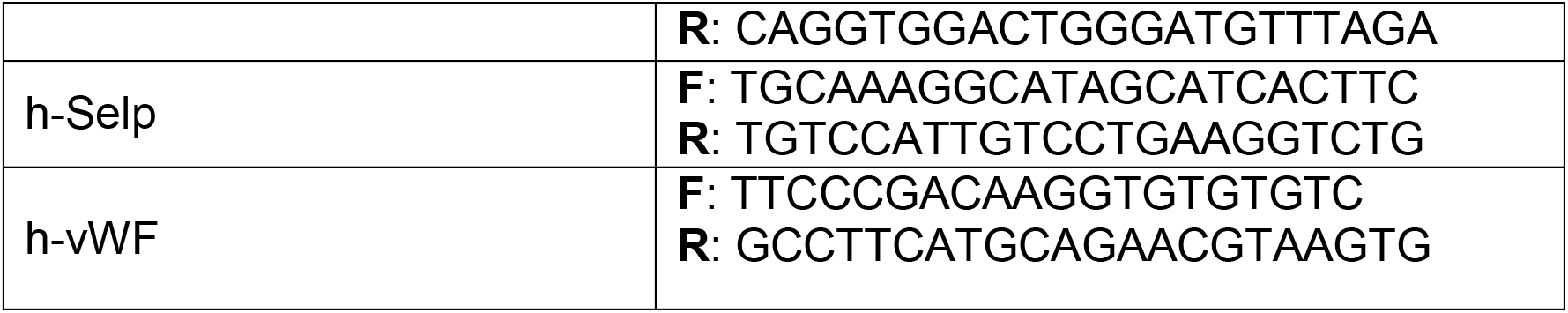
RT-qPCR primers used in this study.

### vWF Strings

HUVECs were plated on Ibidi µ-slide I^0.4^ Leur (IbiTreat) and subjected to HSS, LOSS and static conditions as described previously, with an additional static control. After 48h of flow application, all slides apart from the additional static slide were stimulated with 150 µM histamine for 10 mins at 2.5 dyn/cm^2^ (37°C, 5% CO_2_). The additional static slide was exposed to 1 µM PMA for 10 mins at 2.5 dyn/cm^2^ (37°C, 5% CO_2_), acting as a positive control. After 10 mins of stimulation, each slide was transferred to a new fluidic unit containing 4% PFA, which was applied at 2.5 dyn/cm^2^ for 10 mins at room temperature to fix the cells. Slides were then used for immunofluorescent staining (without permeabilisation) with vWF.

### Immunocytochemistry

After 48 hours of exposure to shear stress, cells were fixed with 4% paraformaldehyde (PFA) for 10 mins at room temperature, washed with PBS and then permeabilised with 0.1% Triton-X solution for 10 mins. Cells were incubated with primary antibody (see table 2) for 1 hour, then washed with PBS and incubated with the relevant species-specific fluorescent dye-conjugated secondary antibody for 30 mins. Cells were washed with PBS and incubated with HOECHST 33342 (Cell signalling) before being mounted with Ibidi mounting medium.

**Table 2:** Antibodies used in Immunocytochemistry.

| Primary Antibody | Species | Application | Working Dilution | Supplier |
| --- | --- | --- | --- | --- |
| Anti-vWF | Mouse | Cells | 1:200 | DAKO |
| Anti-vWF | Rabbit | Aorta | 1:50 | DAKO |
| Anti-GM130 | Mouse | Cells | 1:200 | BD Biosciences |
| Anti-Angpt2 | Goat | Cells | 1:100 | R&D Systems |
| Anti-CD31 | Rat | Aorta | 1:50 | R&D Systems |
| Secondary Antibody |  | Application | Working Dilution | Supplier |
| AlexaFluor 488 anti-mouse IgG |  | Cells | 1:300 | Jackson ImmunoResearch Labs |
| AlexaFluor 488 anti-goat IgG |  | Cells | 1:300 | Jackson ImmunoResearch Labs |
| AlexaFluor 594 anti-mouse IgG |  | Cells | 1:300 | Jackson ImmunoResearch Labs |
| AlexaFluor 594 anti-rabbit IgG |  | Aorta | 1:300 | Jackson ImmunoResearch Labs |
| AlexaFluor 488 anti-rat IgG |  | Aorta | 1:300 | Invitrogen |

### Aorta Extraction and Staining

Animal use was authorised by the University of Leeds Animal Ethics Committee. In order to observe the effect of shear stress on WPBs in vivo, the aortic arch and descending aorta of C57/B mice were flushed with PBS fixed with 4% PFA. Once fixed, aortas were extracted and permeabilised with 0.1% Triton X-100 before blocking for 1 hour. Aortas were then incubated overnight at 4°C with primary antibody (see supplementary table for primary antibodies) followed by incubation with relevant species-specific fluorescent dye-conjugated secondary antibody for 1 hour (see supplementary table for secondary antibodies). The whole aortas (including the arch) were then cut longitudinally to expose the lumen, and mounted on a cover slip, en face, with fluoromount (containing DAPI) before imaging with confocal microscopy.

### Microscopy

All phase contrast imaging of cells on Ibidi slides was performed on the IncuCyte ZOOM (Essen Bioscience). 32 images/ slide were taken in a preset grid.

### Wide-Field Deconvolution Microscopy

Fluorescent microscopy was performed on an Olympus inverted microscope supported by Applied Precision and CellSense deconvolution systems. A z-stack consisting of 10 focal planes at 0.2 µm intervals was taken and iterative deconvolution (5x) was performed using a proprietary algorithm. A maximum intensity projection was applied and further processing and analysis performed on Fiji. All wide-field imaging was performed at room temperature.

### Confocal Microscopy

High-resolution microscopy was performed using a Zeiss LS880 inverted or upright laser-scanning confocal microscope. Images were captured using a 20x air objective for vWF string analysis or 63× oil objective for Golgi analysis. Due to the size of the vWF strings, a tile scan was used (2×2) to increase the capture area. All the images were acquired and processed with Zen software. All confocal imaging was performed at room temperature.

### scRNAseq analysis

scRNAseq FastQ data files generated by Andueza et al. 2020 were obtained from the NCBI BioProject repository (accession number PRJNA646233). Data files were downloaded and stored using the University of Leeds High Performance Computing system – arc3. Arc3 was also used to process the data with Cell Ranger Software, aligning sequencing reads to the mouse reference genome with STAR aligner. H5 files were generated which were then processed using Seurat R package to visualise the data. The code used to analyse the data was kindly made available with the study by Andueza et al. ^26^ (https://github.com/JoLab-Emory/SingleCell). Briefly, a quality control step was performed, removing low quality cells and doublets based on a threshold of unique feature counts, discarding those with less then 200 or more than 7600 feature counts. Cells with lower than 10% of mitochondrial counts were also discarded. Datasets were then merged, normalised, scaled, clustered and visualised by Uniform Manifold Approximation and Projection (UMAP).

### Image Processing and Analysis

Maximum intensity projections were generated using DeltaVision SoftWoRx or Zeiss Zen software and subsequently analysed in Fiji. ImageJ macros were used for the quantification of WPB number and Feret, vWF string length and Golgi fragmentation.

### Analysis of Fluorescent Microscopy Images

Analysis of wide-field deconvolution and confocal images was carried out using automated ImageJ macros (Supplementary macro 1). Briefly, to identify and analyse WPBs, vWF strings and Golgi objects, channels were split and background subtraction applied if necessary. A local threshold was applied to the channel of interest, followed by particle analysis to determine a count, Feret diameter, Feret angle and area of each object. To determine the number of objects or WPBs per cell, the total per field of view (FOV) was divided by the number of nuclei within each image. WPB size in the mouse aortas (n=2) was determined using images (N=8) that were equally enhanced according to the histogram. The red (vWF) channel was set to a minimum auto threshold and the particle analyser in ImageJ was used to measure particles at 0.3 to infinity and circularity 0.05 to 1.

### Statistics

All images were acquired in a random manner and analysis equally automated using Fiji macros and therefore no blinding was necessary. All averaged data are presented as mean ± SEM. One-way ANOVA was performed to determine whether significant differences existed among three or more groups, coupled with Bonferroni post hoc test. Two sample t-test was performed when comparing two sample data groups after normality was approached by a log-transformation, and equal variance confirmed. Significance values are shown in the figures or in figure legends. Statistical significance was considered to exist at probability *p* < 0.05 (### < 10^−15^, *** < 0.001, ** < 0.01, * < 0.05). Where comparisons lacked an asterisk or marked NS, no significant difference between groups was observed. OriginPro 2020 software and R studio was used for data analysis and presentation. *n* = represents the number of independent repeats.

## Results

To mimic conditions of the physiological laminar shear stress sensed by ECs lining the aorta (see Supplementary Table 1 blood vessel type and shear stress) we cultured confluent Human Umbilical Vein Endothelial Cells (HUVECs: a robust model of WPB biogenesis and function) and Human Aortic Endothelial Cells (HAECs: a flow-responsive and physiologically relevant model of arterial ECs) under constant laminar flow at 10 dynes/cm^2^ for 72 hours (high laminar shear stress: HSS). We observed that, as expected, under these conditions cells consistently aligned in the direction of flow and expressed the atheroprotective transcription factor KLF2 (Fig. 1). Compared to HSS, cells cultured under low oscillatory shear (LOSS) or static conditions failed to align in the direction of flow and did not express KLF2 (Fig. 1).

**Figure 1.**
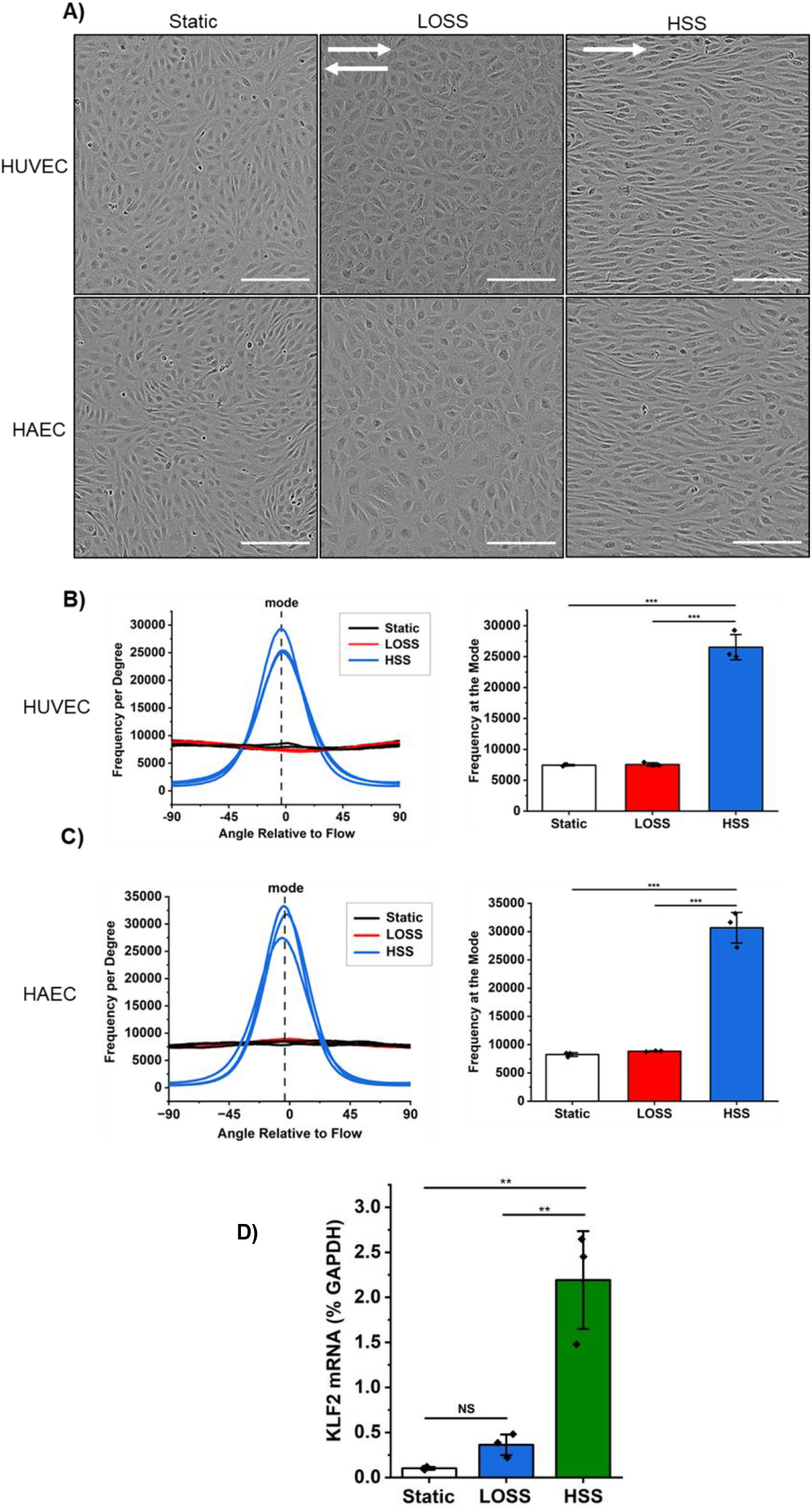
Endothelial cell alignment and KLF2 expression under different flow conditions. A) Representative images from each condition using the IncuCyte ZOOM. Arrows indicate direction of flow. Scale bar = 200 µm. B) and C) Histograms to show the alignment of HUVECs and HAECs relative to the direction of flow and frequency at the mode of each histogram. *n* = 3 for each condition. D) Graph of quantitative PCR change in cycle threshold (ΔCT) analysis (± SEM) of KLF2 mRNA expression in each shear stress condition. Expression was measured as a percentage of the housekeeping gene control (GAPDH). *n* = 3 for each condition. Bars represent mean ± SEM. ** *p* < 0.01 by one-way ANOVA with Bonferroni post-hoc test.

Since cargo released from WPBs evokes thrombi formation, immune responses, vasoconstriction and cell migration, changes in the number, size, and cargo recruitment have the potential to promote cardiovascular disease. WPBs are formed solely due to the production of vWF, thereby an increase in the number of WPBs could increase the thrombotic potential of these cells. Hence, we first explored the effect of HSS and LOSS on the numbers of WPBs generated by ECs using vWF immunostaining as a marker of WPBs together with automated segmentation and analysis (Supplementary macro 1). All experiments were performed in triplicate and ten fields of view captured per biological repeat (approx. 56000 WPBs total per condition). There were no significant differences observed in the numbers of WPBs per cell or the number of cells that contain WPBs, between ECs cultured under HSS or LOSS conditions, (Fig. 2), but we observed less WPBs in HAECs than HUVECs. These data suggest that aberrant shear stress does affect WPB biogenesis in ECs.

**Figure 2.**
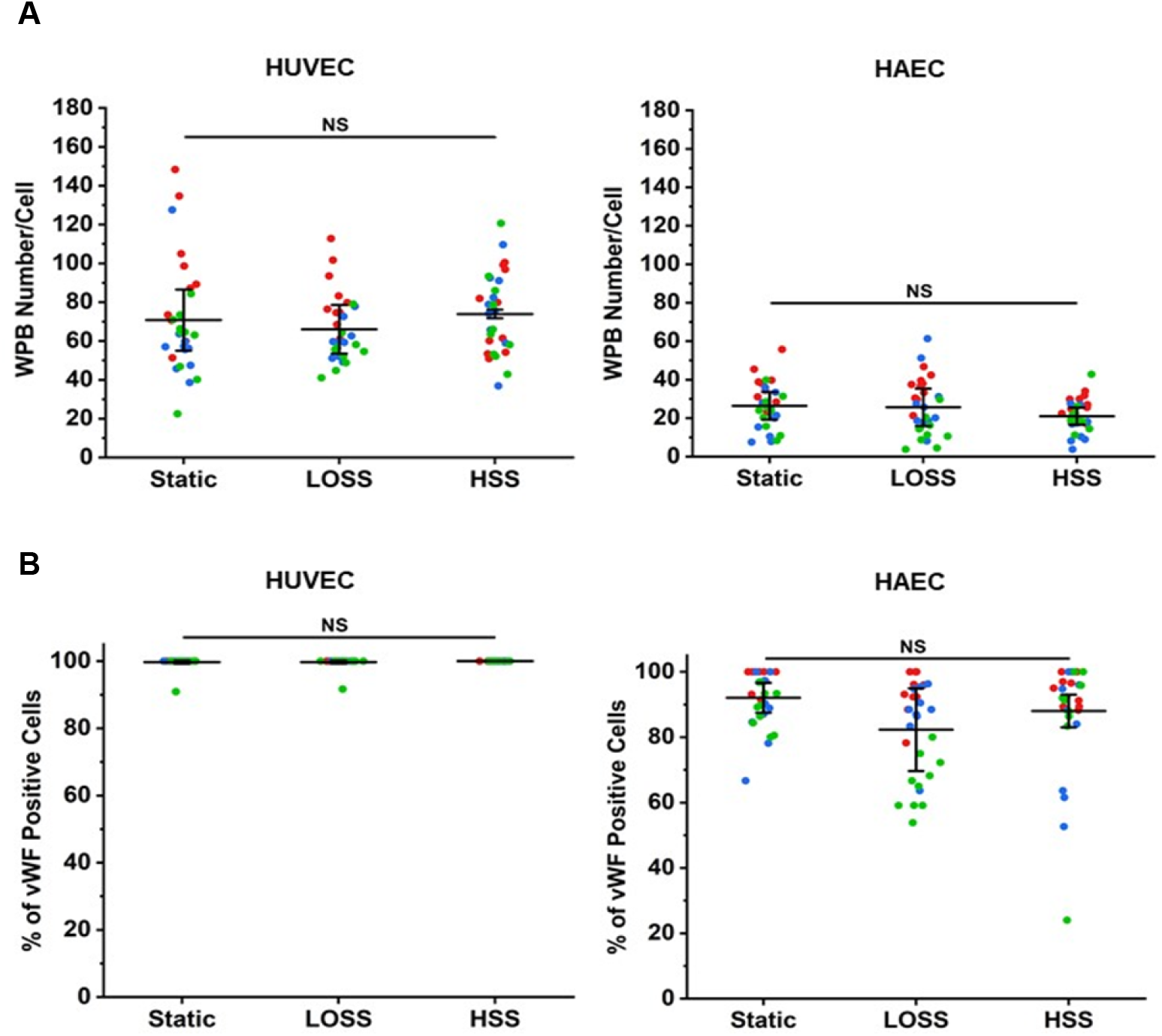
WPB numbers in HUVEC and HAEC do not change under different flow conditions. A) Number of WPBs per cell, based on the total number of cells (nuclei) in each image. B) Percentage of vWF positive cells compared to the total number of cells (nuclei) in each image. All super plots show mean ± SEM. *n* = 3 (red, green, blue represent each independent repeat), NS = not significant by one-way ANOVA with Bonferroni post-hoc test.

Since *in vivo* observations have suggested an impact of blood flow on WPB body size, we questioned if WPBs generated in ECs cultured under HSS are a different size to those cultured under LOSS conditions. We measured the lengths of WPBs using vWF immunostaining as a marker of WPB morphology together with the automated segmentation and analysis (Supplementary macro 1) and normalization to cells cultured under static conditions (as a non-treatment control). WPB length was significantly shorter in HUVECs exposed to HSS (0.81 ± 0.03 μm) compared to WPBs in HUVECs exposed to LOSS (1.08 ± 0.01 μm) or static conditions (1.16 ± 0.02 μm) (Fig 3. A, B, C). The flow-dependent changes in length were also observed in HAECs exposed to HSS (0.84 ± 0.03 μm), LOSS (0.99 ± 0.002 μm) and static conditions (1.17 ± 0.05 μm) (Fig 3. D, E, F). WPBs in HAECs exposed to LOSS are also significantly shorter than WPBs in static HAECs, but not to the same extent as in HSS. Cumulative frequency analysis demonstrating the distribution of WPB lengths under each condition shows a shift to a population of shorter WPBs in HSS conditions compared to LOSS and static in both populations of ECs (Fig 3. C and F).

**Figure 3.**
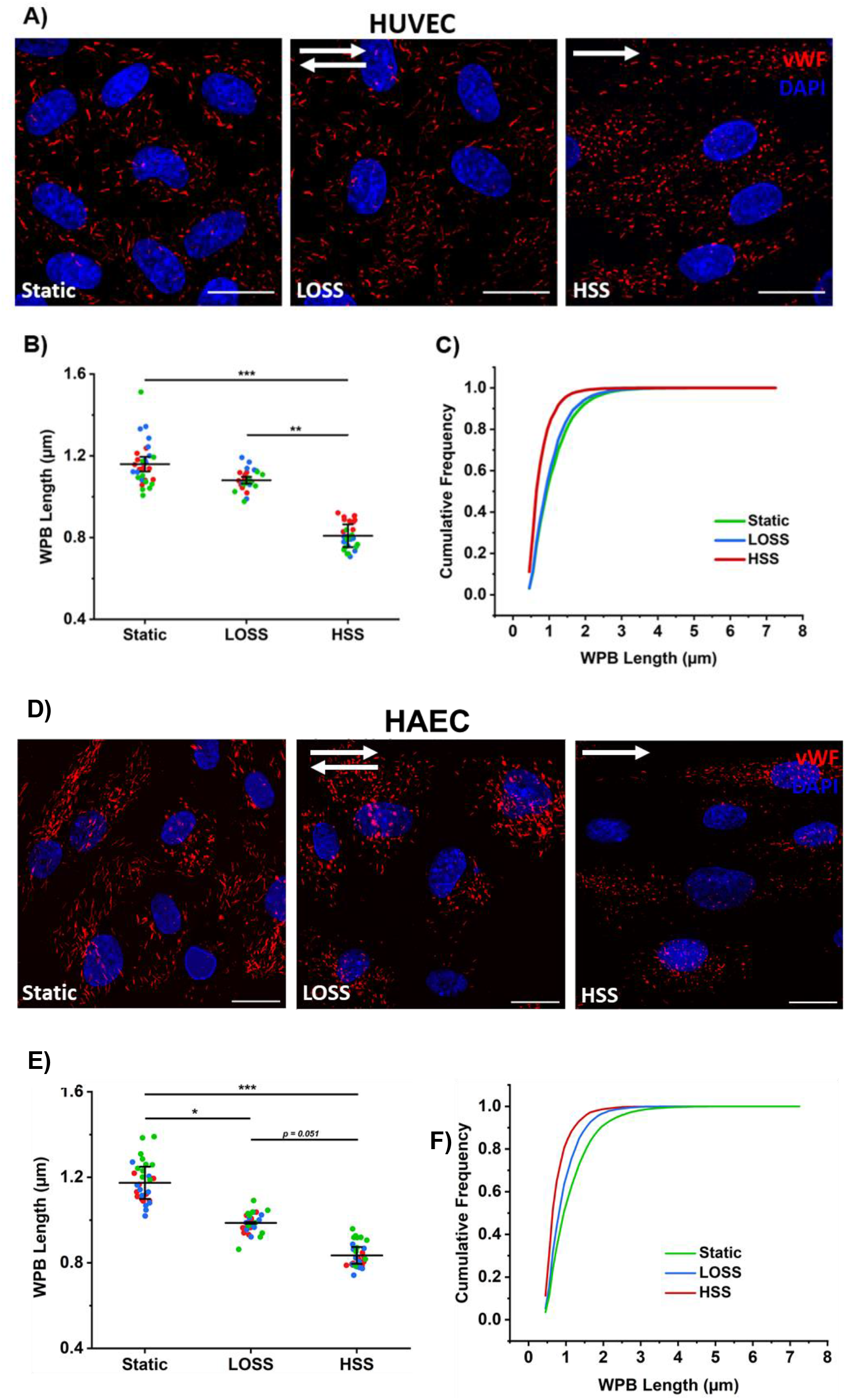
WPB length in HUVEC and HAEC under different flow conditions. A) Representative images (x60 objective) of WPBs (red) in HUVECs from each flow condition. Scale bar = 20 µm. B) Super plot to show mean WPB length ± SEM in each condition, *n* = 3. Each data point represents the mean WPB length from one image (10 images per condition, per repeat). Each colour represents an independent repeat. C) Cumulative frequency graph of WPB length of all WPBs measured (65599). D) Representative images (x60 objective) of WPBs (red) in HAECs from each flow condition. Scale bar = 20 µm. E) Super plot to show mean WPB length ± SEM in each condition, *n* = 3. Each data point represents the mean WPB length from one image (10 images per condition, per repeat). Each colour represents an independent repeat. F) Cumulative frequency graph of WPB length of all WPBs measured (56316). * *p* < 0.05, ** *p* < 0.01, *** *p* < 0.001.

The ECs lining the aorta are subjected to both laminar high shear in straight sections and aberrant shear in areas of bifurcation and around the aortic arch, which ultimately promotes plaque formation. To understand if flow-specific regulation of WPB size in ECs is observed *in vivo*, we imaged vWF in aortas from wild-type C57/black mice. As previously observed in rats ^27^, we observed an increase (0.92 +/- 0.22 vs 0.58 +/- 0.04 µm) in the average size of WPBs (vWF: red) in ECs lining the aortic arch (Fig. 4 A, B), where the ECs are not aligned (Fig. 4 CD31 staining: green), as compared to the straight aorta where ECs align in the direction of flow. These data suggest that the changes in WPB size observed in our *in vitro* model may mimic those observed *in vivo*.

**Figure 4.**
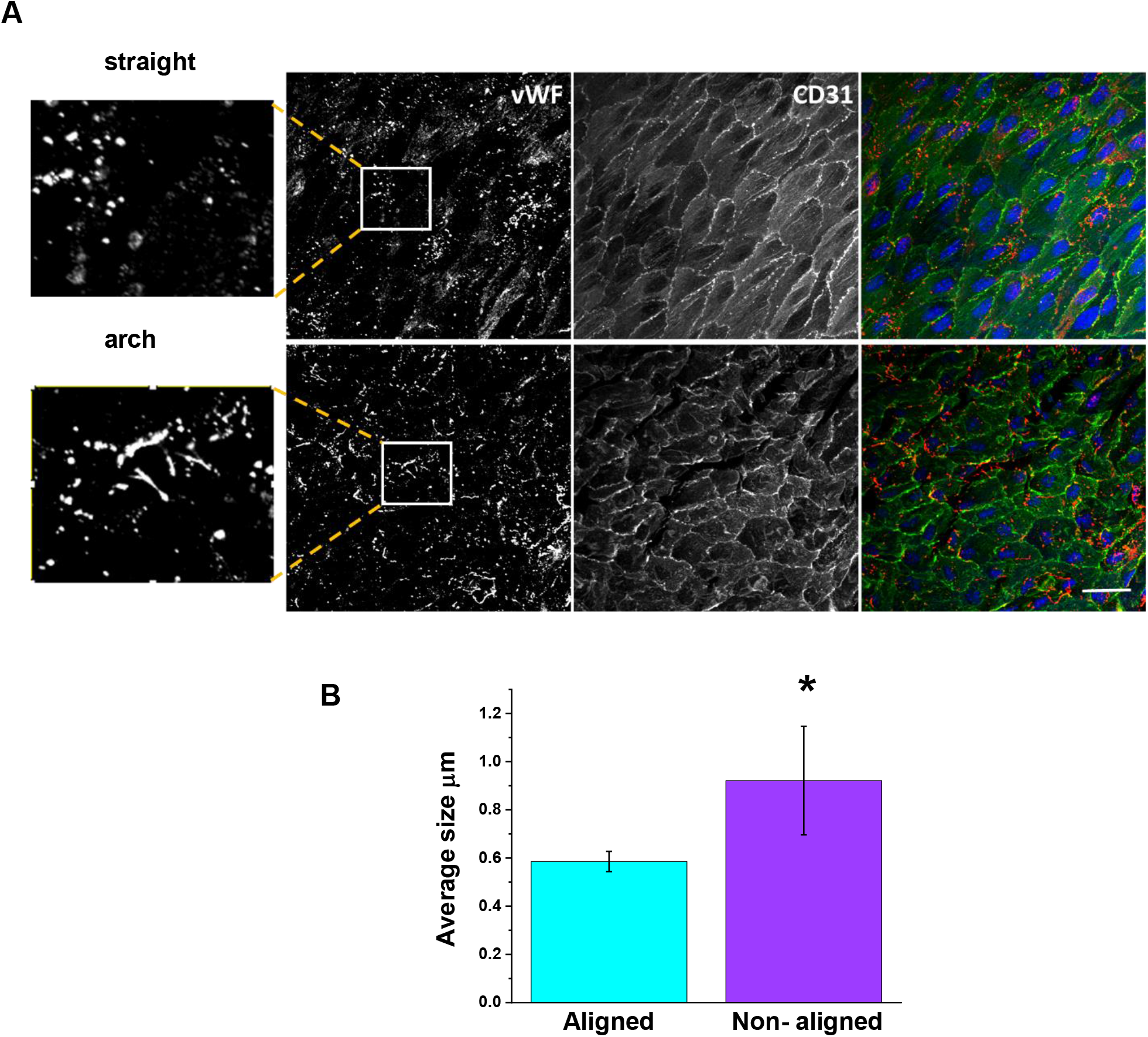
Change in size of WPBs observed in endothelial cells of the mouse aortic arch. A) WPBs (vWF: red) and CD31 (green: cell membrane) were examined by en face immunostaining of ECs lining the mouse aorta. The box depicts the region of interest on the vWF image that is magnified to the left. The CD31 staining demonstrates the alignment of the cells in the arched regions versus the straight regions. Scale bar = 30 µM. B) Mean data ± SD of the average size (µm) of the WPBs in the regions of the aorta where the ECs aligned versus the non-aligned regions. n/N = 2/8. * p < 0.05

**Figure 5.**
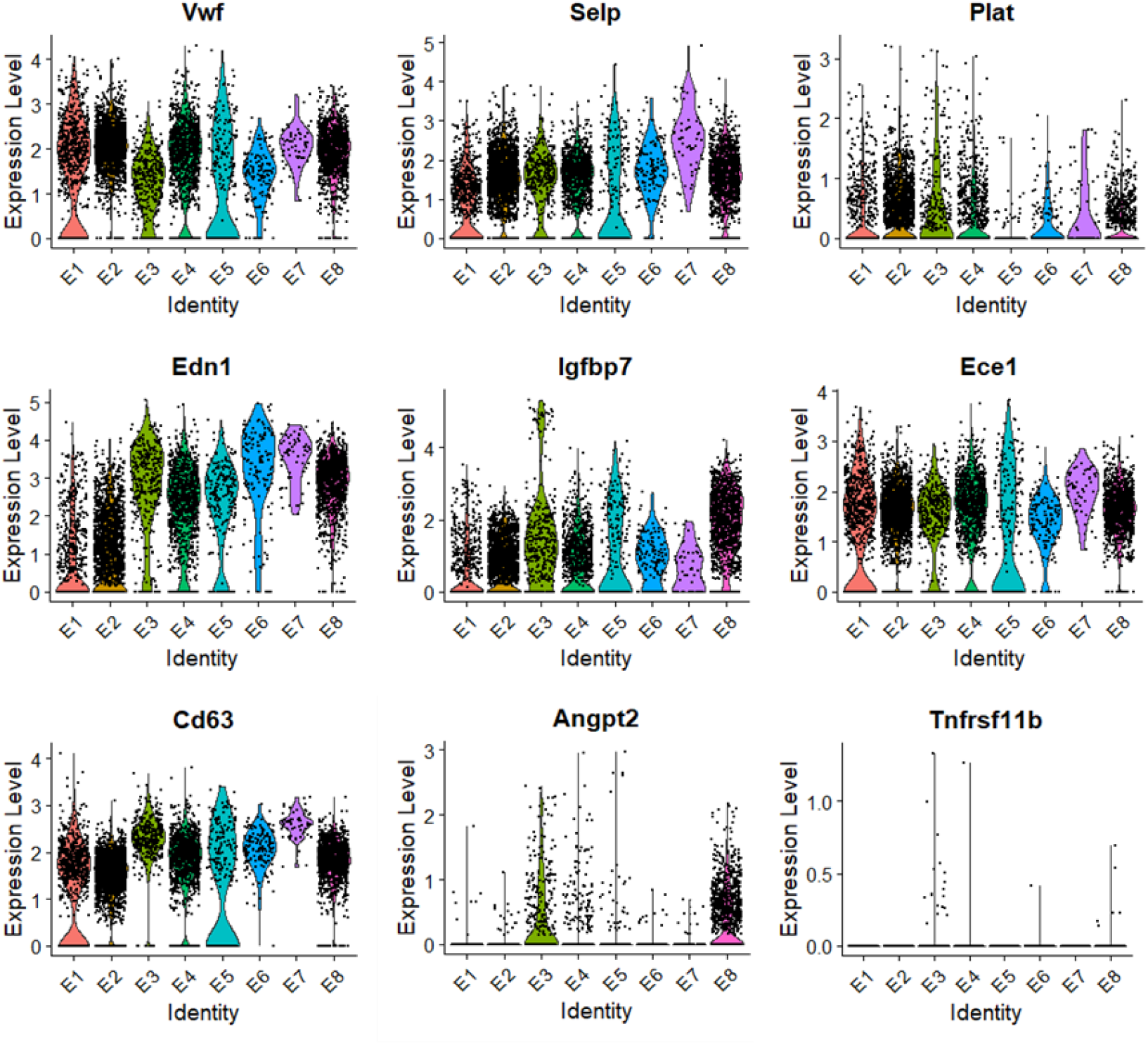
Violin plots to show WPB cargo gene expression. Each of the endothelial clusters were analysed for. expression of WPB cargo listed in Supplementary Table 2.

Changes in WPB length could be mediated by changes in vWF gene expression or changes in how vWF is packaged into the WPBs. Moreover changes in the expression levels of other genes encoding for WPB cargo proteins could impact WPB function. Previous research ^26^ using single cell RNA sequencing (scRNA-seq) from ligated (a model of disturbed laminar flow: d-flow) and non-ligated (a model of high laminar flow: s-flow) carotid arteries show that disturbed flow induces changes in gene expression that result in a transition of ECs from an anti-inflammatory and anti-atherogenic phenotype to a pro-inflammatory and pro-atherogenic phenotype. We utilised this dataset to investigate changes in gene expression of WPB cargo and WPB-associated proteins (Supplementary Table 2). Our analysis of the scRNA-seq data reveals cell phenotypes and EC clusters that concurs with previous studies (Supplementary Fig. 1). Within the clusters, 8 distinctive EC clusters were identified. These were characterised by the size of each cluster in each condition, and using gene markers known to be up and down-regulated in different types of flow (Supplementary Table 3). Based on the cluster size in each flow condition, E1-E4 consisted of ECs mostly exposed to s-flow (2D-R and 2W-R). Clusters E5 and E7 were mostly ECs exposed to acute d-flow (2D-L), whilst E6 and E8 contained cells exposed to chronic d-flow (2W-L). In terms of gene marker expression, cluster E2 had the highest levels of the known mechanosensitive genes (KLF2 and KLF4) and therefore labelled as most representative of ECs in s-flow (i.e. HSS). Cluster E8 had the highest level of thrombospondin-1, a gene known to be upregulated in d-flow, and therefore labelled as the most representative of ECs in chronic d-flow (i.e. LOSS). The expression of WPB cargo were investigated in each EC cluster. The expression of vWF and P-selectin are not significantly different between E2 (s-flow) and E8 (d-flow) clusters, whereas Angpt2 is upregulated in d-flow conditions (as expected) (Figure 55). There is significant differences in the expression of other WPB cargo between E2 and E8 clusters including; endothelin-1 (Edn1), insulin growth factor binding protein 7 (Igfbp7), CD63 and tissue plasminogen activator (Plat) (Fig. 6).

**Figure 6.**
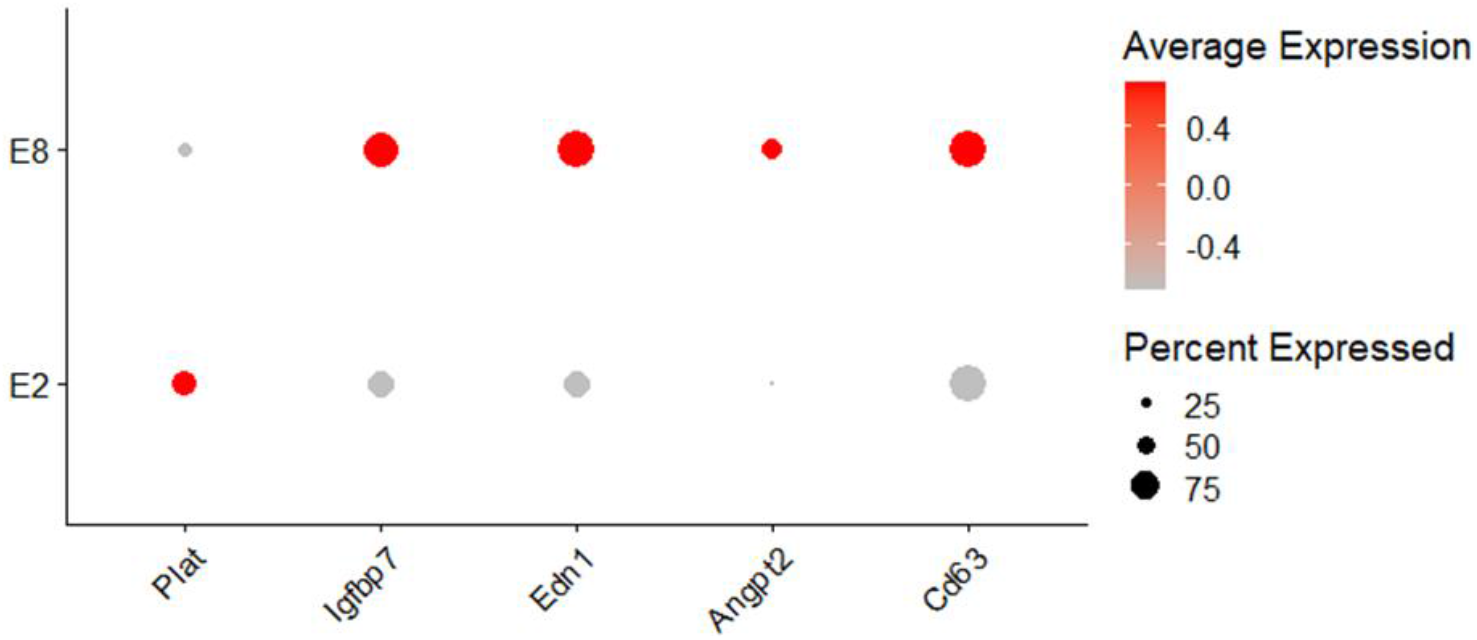
Dot plot to exhibit the average expression per condition and percentage expression of significantly different WPB cargo genes between s-flow (E2) and chronic d-flow (E8). Plat = plasminogen activator; Igfbp7 = Insulin growth factor binding protein7; Edn1 = endothelin1; Angpt2 = Angiopoietin2.

RT-PCR analysis of HUVECs exposed to HSS, LOSS or static conditions for 48h using our *in vitro* model, demonstrated gene expression levels of vWF, P-selectin and Angpt2 that are consistent with the *in vivo* scRNA-seq data (Fig. 7A). The data suggests the *in vitro* model reflects the *in vivo* data, where shear stress conditions have no effect on the expression levels of vWF or P-selectin but LOSS (as a model of d-flow) induces an increase in Angpt2. We questioned if the increase in expression culminated in an increase in recruitment of Angpt2 to WPBs. Immunostaining and imaging of Angpt2 in HUVECs demonstrates that LOSS-dependent increase in gene expression facilitates recruitment of Angpt2 to WPBs (Fig. 7B).

**Figure 7.**
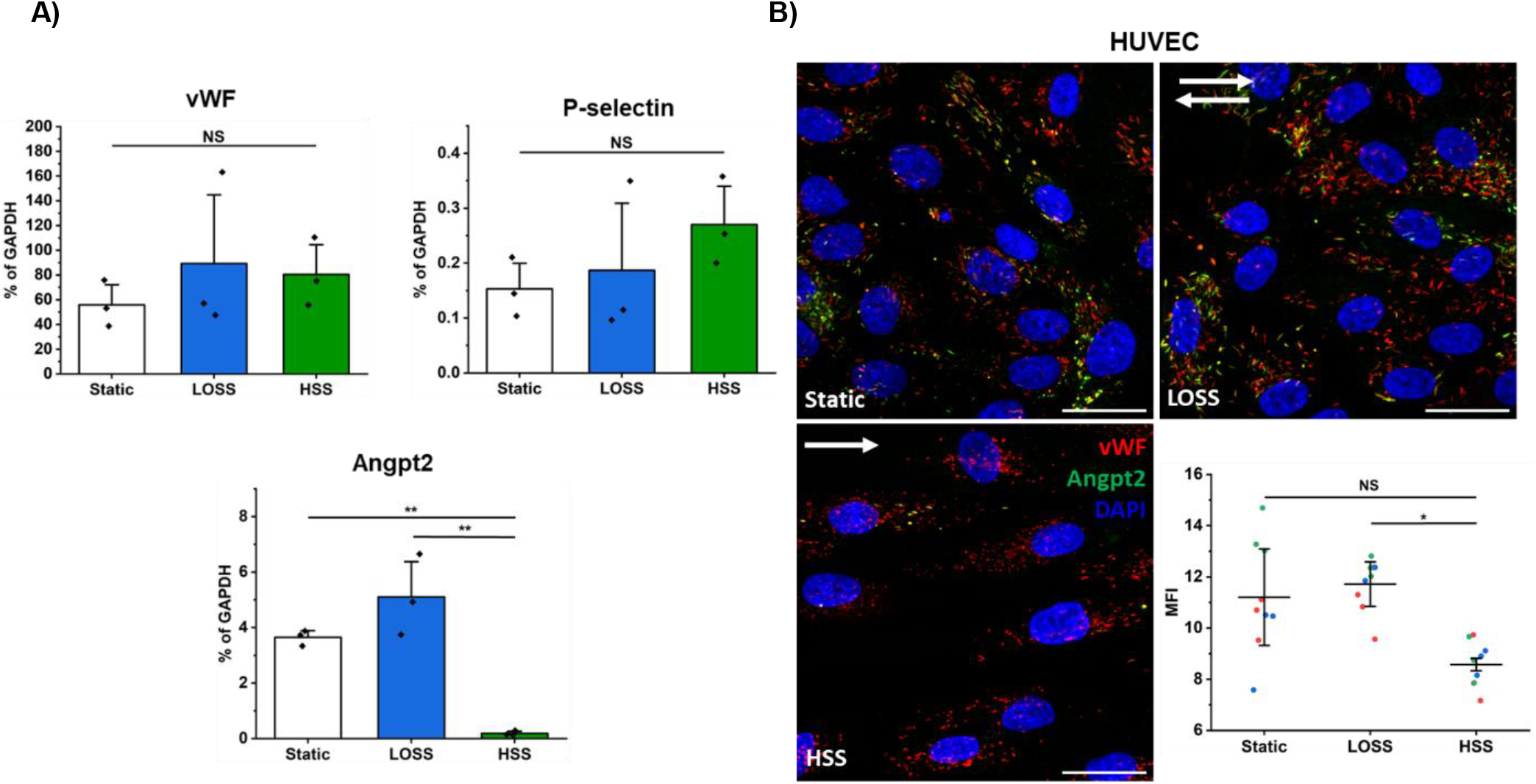
RT-PCR analysis shows expression of genes using the in vitro Ibidi system matched the mouse scRNAseq and flow-regulated Angpt2 is recruited to WPBs. A) Graphs of quantitative PCR change in cycle threshold (ΔCT) analysis (± SEM) of vWF, P-selectin and Angpt2 mRNA expression in cells cultured in static, HSS or LOSS shear stress conditions. Expression was measured as a percentage of the housekeeping control (GAPDH). *n* = 3 for each condition. Bars represent mean ± SEM. B) Immunofluorescent images of Angpt2 (green) and VWF (red: marker of WPBs) in HUVECs under static, HSS or LOSS. Nuclei = DAPI (blue). Arrows indicate direction of flow. Scale bar = 30 µM. The graph is the mean data (MFI: mean fluorescent intensity) of the images depicting Angpt2 in HUVECs. * *p* < 0.05 by one-way ANOVA with Bonferroni post-hoc test.

The data thus far suggests that LOSS regulates WPB size and cargo recruitment, both of which could impact WPB function. The LOSS-evoked increase of Angpt2 expression and the impact on endothelial-mediated angiogenesis has been previously determined ^28^. Thereby here, we focussed on the mechanisms underlying LOSS-evoked changes in WPB size.

Since the shape and size of WPBs are due to the expression and multimerization of vWF ^29 30^, we questioned how LOSS could induce longer WPBs if the expression levels of vWF remain unaffected by flow in the models we analysed. Ferraro et al previously demonstrated that the unit lengths of WPBs correlate with discreet ‘quantas’ or ‘boluses’ of multimeric vWF, and the number of vWF ‘quantas’ that are packaged into the nascent WPB at the Golgi apparatus determines WPB size ^31^. This study further demonstrated that the number of quanta packaged into the emerging WPB is a reflection of the structural arrangement of the Golgi apparatus, where fragmented small Golgi stacks insert single vWF ‘quanta’ into WPBs making them shorter. Conversely, continuous ribbon-like structures allow co-packaging of vWF ‘quanta’, resulting in longer WPB lengths. To understand if LOSS impacts the structure of the Golgi, thereby promoting longer WPBs, we compared the morphology of the Golgi apparatus using immunofluorescent staining and imaging of HUVECs cultured under HSS or LOSS conditions (Fig. 8). LOSS evoked less fragmented (3.17 ± 0.18) Golgi structures as compared to HSS (4.55 ± 0.30). These data indicate that the changes we observe in WPB size could be regulated by flow-dependent changes in Golgi morphology, thereby impacting vWF packaging.

**Figure 8.**
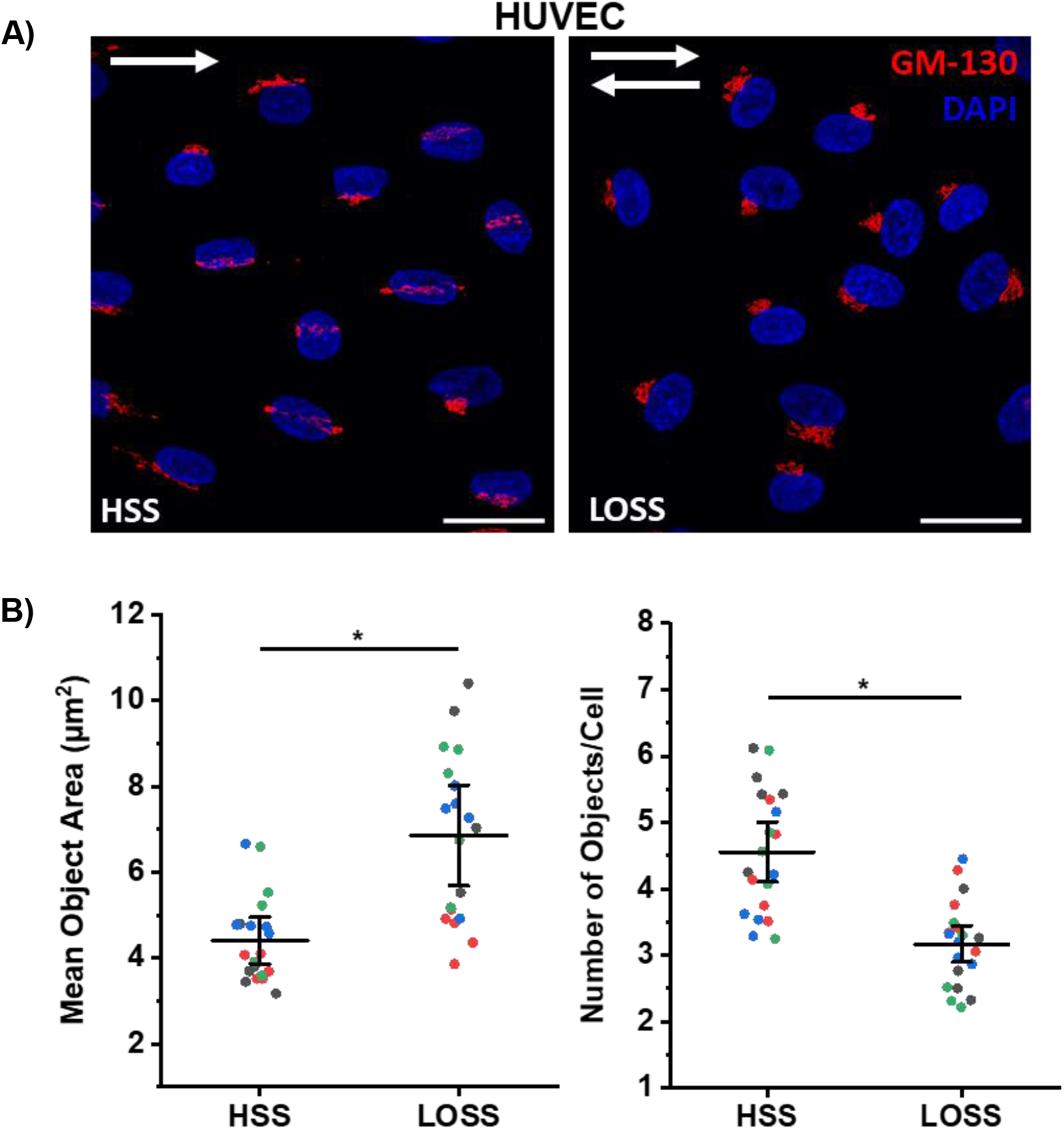
HSS causes increased Golgi fragmentation compared to LOSS. A) Representative images (x60 objective) of Golgi apparatus (red) in HUVECs from each flow condition. Scale bar = 30 µm. B) Super plots to show mean number of Golgi objects identified per cell and the mean area of these objects ± SEM in each condition, *n* = 3/4. Each data point represents the mean value calculated from one image (10 images per condition, per repeat). Each colour represents a separate independent repeat. * *p* < 0.05 by t-test.

The number of vWF quanta packaged into a WPB impacts vWF function, where stimulated cells containing short WPBs release shorter vWF strings resulting in less platelet attachment ^32^. We analysed the effect of flow on vWF string formation (Fig. 9). HUVECs grown under HSS or LOSS for 48 hours were then stimulated with 150 µM histamine at 2.5 dynes/cm^2^ to stimulate WPB exocytosis and the string lengths were analysed. In cells cultured under LOSS the strings were significantly longer (24.98 µm ± 3.41) compared to those under HSS conditions (10.96 µm ± 0.57). Together these results indicate that LOSS elongates WPBs, by controlling the morphology of extended Golgi stacks that allow for loading of multiple VWF quanta into newly forming WPBs.

**Figure 9.**
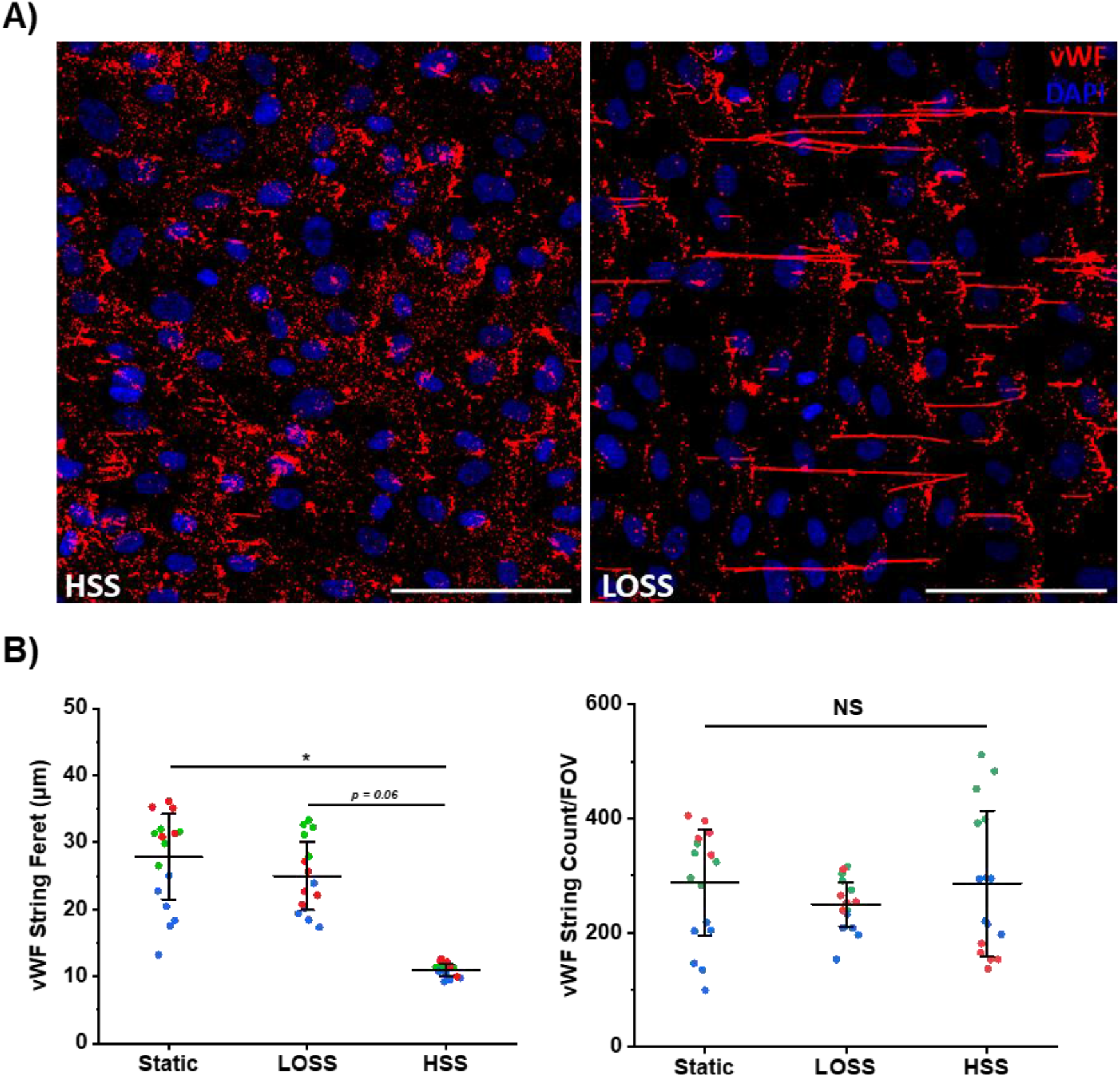
Exposure to HSS results in shorter vWF strings upon histamine stimulation compared to LOSS and static conditions. A) Representative images (x20 objective, 2×2 tile scan) of vWF strings (red) from HUVECs with 10 minute histamine (or PMA where stated) stimulation after 48h exposure to each flow condition. Scale bar = 100 µm. B) Super plots show mean Feret diameter of vWF strings and the number of vWF strings per FOV ± SEM in each condition, *n* = 3. Each data point represents the mean value calculated from one image (5 images per condition, per repeat). Each colour point represents a separate independent repeat. * *p* < 0.05 by t-test.

## Discussion

WPBs are generated by the expression and multimerization of vWF but their vital role in homeostasis is supported by the recruitment of other functionally distinct cargo (review see ^33^). To generate a stimuli-specific response the differential release of cargo can be regulated by sorting mechanisms at the plasma membrane ^22 21 19^, however, it is also becoming increasingly clear that ECs respond to their environment by generating subpopulations of WPBs, carrying specific cargoes, that are trafficked in a context-dependent manner ^23 24 25^. ECs are constantly influenced by the pressures exerted by the flow of the blood including hydrostatic forces, tension and shear stress ^34^ where shear stress, the force exerted on the apical surface of ECs, is dominant ^35^. Studies have shown that areas of aberrant or disturbed shear evoked by oscillatory flow, such as occurs at vessel bifurcations and arched regions, are susceptible to plaque formation. Since atheroprone regions tend to have raised plasma levels of the proinflammatory and prothrombotic mediators stored in WPBs, such as vWF, P-selectin and Angpt2 it is reasonable to ask how shear stress impacts WPBs. Therefore we explored if LOSS, as a model for regions of disturbed flow, evoked differences in the number or size of WPBs as compared to WPBs stored in ECs cultured under physiological HSS.

Using a validated Ibidi flow system, we determined that LOSS induces the generation of significantly longer WPBs as compared to WPBs in cells cultured under HSS conditions. This was consistent in both HUVECs, ECs that act as a model of robust WPB biogenesis and function, and HAECs, ECs that are subjected to changes in haemostatic pressures. scRNA-seq data taken from a mouse model of aberrant flow, suggests that disturbed flow impacts the expression levels of some WPB cargo such as Angpt2, a known flow-responsive factor that promotes angiogenesis under oscillatory flow conditions ^28^. Here we show that the flow-dependent increase in Angpt2 expression facilitates its recruitment to WPBs. However, an increase in recruitment of these small cargo molecules is unlikely to have a significant effect on WPB size. The length of WPBs has previously been shown to be impacted by the number of ‘boluses’ or ‘quanta’ of multimerized vWF that is inserted into the newly forming WPBs at the site of the Golgi ^31 36 37^. Since no differences in the expression levels of vWF were observed in ECs under LOSS or HSS, which was consistent with scRNA-seq data from the *in vivo* model, we tested if shear stress impacted vWF packaging by measuring the length of the extracellular vWF strings released by the cells after histamine stimulation. The functional consequence of altered WPB size in ECs cultured under LOSS was the secretion of significantly long extracellular vWF strings. Does the differential packaging of vWF into nascent WPBs promote the prothrombotic environment necessary for plaque formation? Long vWF strings show an increased haemostatic potential ^32^. We know that platelet aggregation is readily seen at sites of vascular injury ^38^ and most of this is via vWF released from ECs. Decreased clearance of vWF strings in mice depleted of ADAMTS13 leads to increased platelet aggregation in mesenteric vessels ^39^. Moreover, ADAMTS13 inhibition promotes thrombi formation ^40^ after FeCl_3_ injury and reduces embolization ^39^. It is reasonable to propose that longer vWF strings due to increased bolus packaging, induce increased platelet aggregation in vivo. Our data show that in the aorta WPBs are longer around areas of unaligned regions like the arch which is an atheroprone site. The ex vivo studies require robust measurements of WPB strings in these distinct regions, and further structural observations of vWF formation using EM microscopy.

What are the mechanisms underlying flow-dependent packaging of vWF ‘quanta’? Studies by Ferraro et al. suggest that the structure of the Golgi apparatus influences the size of WPBs ^31^ where a ribbon-like structure permits insertion of more vWF quanta into nascent WPBs than shorter stacks and therefore produces longer vWF strings. Endothelial planar polarity develops in response to flow ^41^ and, although dependent on experimental design and the levels of shear stress, under laminar shear stresses > 7.2 dynes/ cm^2^ there is a bias for the Golgi to align upstream of the nucleus ^42^. However, flow-dependent changes in Golgi morphology has not been widely studied. A study using a microfluidic device that mimics wall shear stress gradients revealed that the projected area of the Golgi was significantly reduced by 51% after flow compared to static conditions ^43^. Whether flow induced fragmentation of the Golgi reflects the packaging of WPBs remains to be established. Our initial studies indicate that HSS induces a more fragmented Golgi suggesting these Golgi stacks regulate packaging of single vWF quanta with the potential to be less thrombotic, therefore the Golgi could be a site of therapeutic intervention. However, there must be a balance between ribbon formation and short fragmented stacks, since, although disassembly of the Golgi ribbon leading to fragmentation is associated with physiological processes such as mitosis, cell division and apoptosis, further fragmented Golgi are also found in pathological conditions such as in neural cells and Alzheimer’s ^44^ and cardiomyopathies ^45^. Indeed a new study by Meli et al ^46^ suggests that ECs isolated from explanted hearts of patients with heart failure due to dilated cardiomyopathy display small rounded WPBs compared to healthy counterparts, due to disordered arrangement of VWF tubules in nascent WPBs emerging from the trans-Golgi network. Rather than an advantage the authors noted that the failure of secreted vWF to form long extracellular strings under flow conditions might allow ECs, in response to stressors such as hypoxia, to adapt to reduce the haemostatic potential of the secreted vWF and hence the likelihood of inappropriate coagulation. Further studies using pathophysiological in vivo models are needed to determine the consequence of altering WPB biogenesis.

One of the observations of this study is that static conditions mimic LOSS. WPB size and string length are significantly increased under static conditions as compared to HSS. Moreover we have shown that cargo such as Angpt2 which is upregulated in cells under LOSS is also expressed in cells cultured under static conditions and recruited to WPBs. We must therefore carefully consider the impact of culturing ECs under static conditions when exploring WPB biogenesis and function.

The advantages of using the Ibidi flow system are the ability to reproducibly dictate the shear stress to which the cells were exposed thus mimicking a more physiological environment and it is an enclosed system so self-contained circulation of media through pumping platforms prevents the removal of soluble factors secreted by ECs that modulate cell–cell communication. However, the cells are grown as a 2D monolayer. To be fully representative of the changing dynamics of a blood vessel there are great efforts to create in vitro models that mimic the human vascular system. The creation of organ-on-a-chip is a great advance, and future technological developments may further focus on replicating cell heterogeneity and the complex three-dimensional structure of tissues in vivo.

## Acknowledgements

This research was supported by a BHF PhD studentship (FS/18/61/34182) as part of a 4yr BHF PhD programme awarded to Ashley Money. We’d like to thank the University of Leeds, Faculty of Biological Sciences Bioimaging facility for their advice (https://biologicalsciences.leeds.ac.uk/facilities/doc/bio-imaging-flow-cytometry).

## Author Contributions

AM and HT performed the experiments. HJ provided the mouse model data. AM, DJB and LM analysed the data and created the tables. LM conceived the study and co-wrote the article with input from all authors. LM generated research funds and supervised the PhD studentship.

## Supplementary Macro 1

**ImageJ WPB count and ferret size macro**

imageTitle=getTitle();//returns a string with the image title

run(“Split Channels”);

selectWindow(“C1-”+imageTitle)

close();

selectWindow(“C2-”+imageTitle);

setOption(“ScaleConversions”, true);

run(“8-bit”);

run(“Subtract Background…”, “rolling=3 sliding”);

run(“Auto Local Threshold”, “method=Bernsen radius=15 parameter_1=0 parameter_2=0 white”);

run(“Analyze Particles…”, “size=0.1-3.00 show=Masks clear add”);

run(“Set Measurements…”, “area perimeter feret’s redirect=None decimal=9”);

roiManager(“Measure”);

saveAs(“Results”, “C:\\Users\\umamo\\OneDrive - University of Leeds\\4-Year PhD

Project\\Flow Experiments\\”+imageTitle+”.txt“);

selectWindow(“C2-”+imageTitle);

close();

selectWindow(“Mask of C2-”+imageTitle);

close();

close(“Results”);

close(“ROI Manager”);

## Supplementary Figures

**Figure S1.**
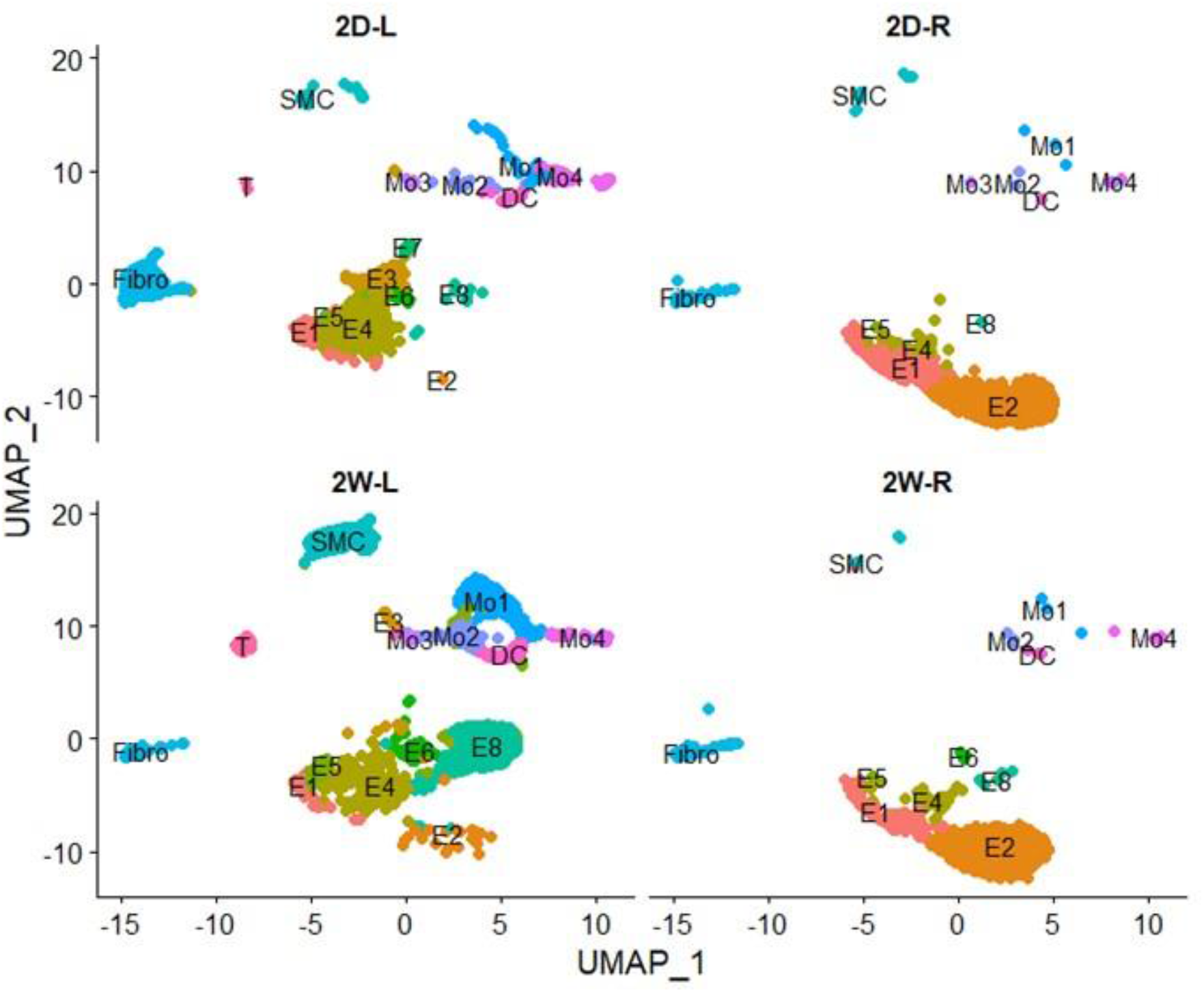
UMAP displaying the clusters of cells in each condition.

## Supplementary Tables

**Table S1.**
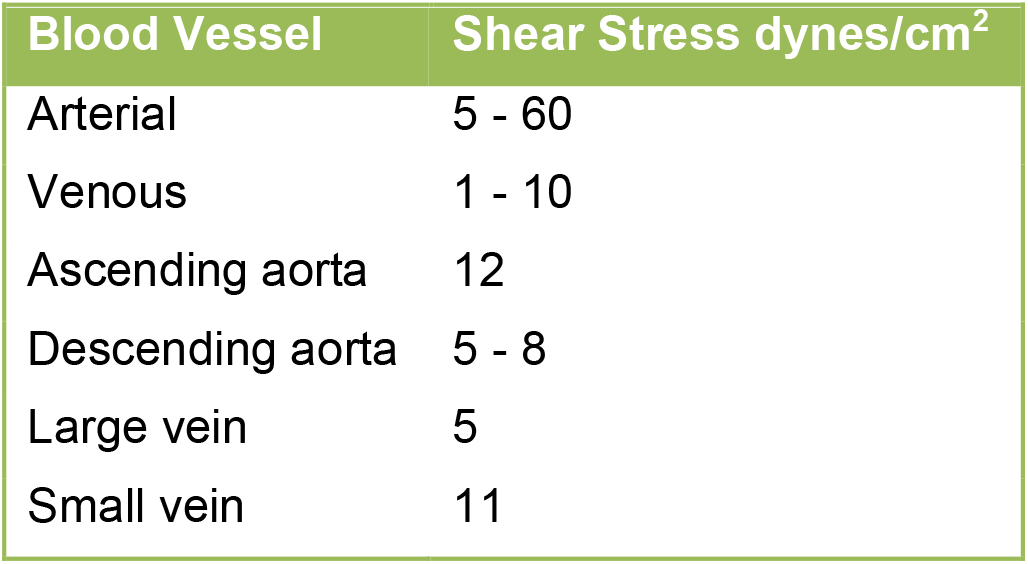
Blood vessel shear stresses. Approximate values taken from McDonald 1974 ^47^Papaioannou et al. 2006 ^48^

**Table S3.** Characterisation of endothelial cell clusters. UMAP identified 8 endothelial cell clusters which were characterised based on gene marker expression and appearance/disappearance of distinct clusters after exposure to different flow types.

| Endothelial Cell Cluster | Characterisation of Flow |
| --- | --- |
| E1 | S-flow (low sensitivity to flow) |
| E2 | Highest levels of known mechanosensitive genes (KLF2/KLF4) |
| E3 | Second highest levels of known mechanosensitive genes (KLF2/KLF4) |
| E4 | S-flow (low sensitivity to flow) |
| E5 | Acute d-flow |
| E6 | Chronic d-flow |
| E7 | Acute d-flow |
| E8 | Chronic d-flow (highest level of Thsp1) |

