## Supplemental Table 2 for "Low oscillatory shear stress regulates Weibel-Palade body size and vWF release"

| wpb_cargo_protein | gene_short_name | reference |
|---|---|---|
| Alpha-2-HS-glycoprotein | AHSG | |
| Angiopoietin-2 | ANGPT2 | Fiedler et al., 2004 |
| Angiopoietin-2 | ANGPT2 | |
| Biglycan | BGN | |
| Calcitonin gene-related peptide | CALCA | |
| Calreticulin | CALR | |
| Monocyte chemoattractant protein- | CCL2 | |
| Eotaxin-3 | CCL26 | |
| CD63 | CD63 | |
| Clusterin | CLU | |
| Collagen alpha-1(I) chain | COL1A1 | |
| Collagen alpha-1(III) chain | COL3A1 | |
| Plasma glutamate carboxypeptidas | CPQ | |
| C-X-C motif chemokine ligand 1 | CXCL1 | |
| Interleukin-8 | CXCL8 | |
| Endothelin converting enzyme 1 | ECE1 | |
| Endothelin-1 | EDN1 | |
| EGF-containing fibulin-like extrace | EFEMP1 | |
| Plasma alpha-L-fucosidase | FUCA2 | |
| Fucosyltransferase 6 | FUT6 | |
| Endoplasim | HSP90B1 | |
| 78 kDa-regulated protein | HSPA5 | |
| Insulin-like growth factor-binding p | IGFBP7 | new confirmed WPB component |
| Insulin-like growth factor-binding p | IGFBP7 | |
| Interleukin 6 | IL6 | |
| Insulin receptor-related protein | INSRR | |
| Integrin alpha-5 | ITGA5 | |
| Lysozyme g-like protein 2 | LYG2 | |
| Epididymis-specific alpha-mannosi | MAN2B2 | |
| Cell Surface glycoprotein MUC18 | MCAM | |
| Matrix Gla protein | MGP | |
| Multimerin-1 | MMRN1 | |
| Puromycin-sensitive aminopeptida | NPEPPSL1 | |
| Nucleobindin-1 | NUCB1 | |
| Protein disulfide-isomerase | P4HB | |
| Protein disulfide-isomerase A3 | PDIA3 | |
| Protein disulfide-isomerase A4 | PDIA4 | |
| Platelet endothelial cell adhesion n | PECAM1 | |
| Tissue Plasminogen Activator | PLAT | |
| Plexin-D1 | PLXND1 | |
| Endothelial protein C receptor | PROCR | |
| Pentraxin-related protein PTX3 | PTX3 | |
| P-selectin | SELP | Bonfanti et al., 1989 and McEver et al., 1989 |
| P-selectin | SELP | |
| Plasminogen activator inhibitor 1 | SERPINE1 | |
| Serpin H1 | SERPINH1 | |
| SPARC | SPARC | |
| Thrombospondin-1 | THBS1 | |
| Osteoprotegerin | TNFRSF11B | |
| V-set and immunoglobulin domain- | VSIG8 | |
| von Willebrand factor A domain-co | VWA5B1 | |
| von Willebrand factor | VWF | Wagner et al., 1982 |
| von Willebrand Factor | VWF | |

| Bona Fide WPB Cargoes (from F | Gene name | References in paper |
|---|---|---|
| vWF | VWF | 6, 7 |
| P-selectin | SELP | 17, 18 |
| Interleukin-8 | CXCL8 | 19, 20 |
| Eotaxin-3 | CCL26 | 21 |
| Calcitonin gene related peptide | CGRP/CALCA | 22 |
| Endothelin | EDN1 | 22, 23 |
| Endothelin converting enzyme | ECE1 | 24 |
| CD63/lamp3 | CD63 | 25, 26 |
| a1,3-fucosyltransferase VI | FUT6 | 27 |
| Tissue-type plasminogen activator | PLAT | 28 |
| Angiopoietin-2 | ANGPT2 | 29, 30 |
| Osteoprotegerin | OPG/TNFRSF11B | 31 |

| wpb_associated_protein/exocyto | gene_short_name | reference |
|---|---|---|
| Golgi resident protein GCP60 | ACBD3 | |
| Lysosomal acid phosphatase | ACP2 | |
| Long-chain-fatty-acid--CoA ligase 4 | ACSL4 | |
| Beta-actin-like protein 2 | ACTBL2 | |
| Actin, gamma-enteric smooth musc | ACTG2 | |
| Alpha-actinin-1 | ACTN1 | |
| Alpha-actinin | ACTN1 | |
| Apha-actinin-4 | ACTN4 | |
| Actin-related protein 3B | ACTR3B | |
| Disintegrin and metalloproteinase d | ADAM9 | |
| Adenylosuccinate synthetase isozy | ADSS2 | |
| Alpha-2-HS-glycoprotein | AHSG | |
| A-kinase anchor protein 12 | AKAP12 | |

| | | |
|---|---|---|
| Retinal dehydrogenase 1 | ALDH1A1 | |
| 10-formyltetrahydrofolate dehydrog | ALDH1L1 | |
| Fructose-bisphosphate aldolase A | ALDOA | |
| Fructose-bisphosphate aldolase C | ALDOC | |
| Angiopoietin-2 | ANGPT2 | |
| Aminopeptidase N | ANPEP | |
| Annexin A1 | ANXA1 | |
| Annexin A2 | ANXA2 | |
| Annexin A2 | ANXA2 | |
| Annexin A3 | ANXA3 | |
| Annexin A4 | ANXA4 | |
| Annexin A5 | ANXA5 | |
| Annexin A6 | ANXA6 | |
| Amyloid beta A4 precursor protein- | APBB2 | |
| Amyloid-like protein 2 | APLP2 | |
| Amyloid beta A4 protein;N-APP;So | APP | |
| ADP-ribosylation factor 1;ADP-ribo | ARF1;ARF3 | |
| ADP-ribosylation factor 5 | ARF5 | |
| Brefeldin A-inhibited guanine nucle | ARFGEF1 | |
| Arfaptin-2 | ARFIP2 | |
| Rho GTPase-activating protein 1 | ARHGAP1 | |
| Rho guanine nucleotide exchange | ARHGEF1 | |
| ADP-ribosylation factor-like protein | ARL1 | |
| PRA1 family protein 3 | ARL6IP5 | |
| Arylsulfatase A | ARSA | |
| Arylsufratase B | ARSB | |
| Acid ceramidase | ASAH1 | |
| ATPase family AAA domain-contai | ATAD3A | |
| Atlastin-3 | ATL3 | |
| Sodium/potassium-transporting AT | ATP1A1 | |
| Sodium/potassium-transporting AT | ATP1A3 | |
| Calcium-transporting ATPase type | ATP2C1 | |
| V-type proton ATPase 116 kDa sul | ATP6V0A1 | |
| V-type proton ATPase 116 kDa sul | ATP6V0A2 | |
| V-type proton ATPase subunit d 1 | ATP6V0D1 | |
| V-type proton ATPase subunit d 1 | ATP6V0D1 | |
| V-type proton ATPase catalytic sub | ATP6V1A | |
| Probable phospholipid-transporting | ATP9A | |
| Ancient ubiquitous protein 1 | AUP1 | |
| BAG family molecular chaperone r | BAG5 | |
| Brain acid soluble protein 1 | BASP1 | |
| B-cell receptor-associated protein : | BCAP31 | |
| BET1 homolog | BET1 | |
| BET1-like protein | BET1L | |
| Biglycan | BGN | |
| Golgin-45 | BLZF1 | |
| Vesicle transport protein SEC20 | BNIP1 | |
| Cell division cycle 42 | C42 | |
| CAD protein | CAD | |
| Caldesmon | CALD1 | |
| Calreticulin | CALR | |
| Coiled-coil domain-containing prote | CCDC186 | |
| T-complex protein 1 subunit gamm | CCT3 | |
| T-complex protein 1 subunit epsilo | CCT5 | |
| T-complex protein 1 subunit theta | CCT6A | |
| T-complex protein 1 subunit zeta | CCT6A | |
| CD63 antigen | CD63 | Vischer UM et al. 1993 |
| CD63 antigen | CD63 | Vischer et al. 1983 |
| Hsp90 co-chaperone Cdc37 | CDC37 | |
| Centrosomal protein of 170 kDa | CEP170 | |
| Clathrin heavy chain 1 | CLTC | Zenner et al. 2007 |
| Clathrin heavy chain 1 | CLTC | |
| Clusterin | CLU | |
| Cytosolic non-specific dipeptidase | CNDP2 | |
| Conserved oligomeric Golgi compl | COG3 | |
| Conserved oligomeric Golgi compl | COG5 | |
| Conserved oligomeric Golgi compl | COG6 | |
| Conserved oligomeric Golgi compl | COG7 | |
| Collagen alpha-1((I) chain | COL1A1 | |
| Collagen alpha-1(III) chain | COL3A1 | |
| Coatomer subunit alpha | COPA | |
| Coatomer subunit epsilon | COPE | |
| Carboxypeptidase D | CPD | |
| Plasma glutamate carboxypeptidas | CPQ | |
| Cleavage and polyadenylation spe | CPSF1 | |
| EF-hand calcium-binding domain-c | CRACR2A | Miteva KT et al. 2019 |
| Exportin-2 | CSE1L | |
| Cathepsin Z | CTSZ | |
| NADH-cytochrome b5 reductase 3 | CYB5R3 | |
| Cytoplasmic FMR1-interacting prot | CYFIP1 | |
| Derlin-2 | DERL2 | |
| 7-dehydrocholesterol reductase | DHCR7 | |
| DnaJ homolog subfamily A membe | DNAJA2 | |

| | | |
|---|---|---|
| DnaJ homolog subfamily C membe | DNAJC5 | |
| Protein dopey-2 | DOPEY2 | |
| Dipeptidyl peptidase 4 | DPP4 | |
| Dipeptidyl peptidase 2 | DPP7 | |
| Desmoglein-1 | DSG1 | |
| Dymeclin | DYM | |
| Cytoplasmic dynein 1 heavy chain | DYNC1H1 | |
| Endothelin-converting enzyme 1 | ECE1 | Russell Fraser D et al. 1998 |
| Elongation factor 1-alpha 1 | EEF1A1 | |
| Elongation factor 1-gamma | EEF1G | |
| Elongation factor 2 | EEF2 | |
| EGF-containing fibulin-like extrace | EFEMP1 | |
| Erythrocyte initiation factor 4A-I | EIF4A1 | |
| Emerin | EMD | |
| Alpha-enolase | ENO1 | |
| Extended synaptotagmin-1 | ESYT1 | |
| Exocyst complex component 6 | EXOC6 | |
| Protein NOXP20 | FAM114A1 | |
| Protein FAM114A2 | FAM114A2 | |
| Protein FAM134C | FAM134C | |
| Protein FAM219A | FAM219A | |
| Fatty acid synthase | FASN | |
| Fumarate hydratase, mitochondria | FH | |
| Four and a half LIM domains prote | FHL2 | |
| Filaggrin-2 | FLG2 | |
| Filamin-A | FLNA | |
| Filamin-A | FLNA | |
| Filamin-B | FLNB | |
| Filamin-B | FLNB | |
| Flotillin-1 | FLOT1 | |
| Flotillin-2 | FLOT2 | |
| Fibronectin | FN1 | |
| Fascin | FSCN1 | |
| Tissue alpha-L-fucosidase | FUCA1 | |
| Plasma alpha-L-fucosidase | FUCA2 | |
| Lysosomal alpha-glucosidase | GAA | |
| N-acetylgalactosamine-6-sulfatase | GALNS | |
| Glyceraldehyde-3-phosphate dehy | GAPDH | |
| Glucosylceramidase | GBA | |
| Golgi-specific brefeldin A-resistanc | GBF1 | Lopes-da-Silva M et al. 2019 |
| GRIP and coiled-coil domain-conta | GCC2 | |
| Rab GDP dissociation inhibitor bet | GDI2 | |
| Glycerophosphodiester phosphodi | GDPD5 | |
| Glycerophosphodiester phosphodi | GDPD5 | |
| GDP-L-fucose synthase | GFUS | |
| Gamma-glutamylcyclotransferase | GGCT | |
| Gamma-glutamyl hydrolase | GGH | |
| GTPase IMAP family member 1 | GIMAP1 | |
| GTPase IMAP family member 7 | GIMAP7 | |
| Gap junction alpha-1 protein | GJA1 | |
| Alpha-galactosidase A | GLA | |
| Beta-galactosidase | GLB1 | |
| Guanine nucleotide-binding protein | GNAI2 | |
| Guanine nucleotide-binding protein | GNAI3 | |
| Guanine nucleotide-binding protein | GNAS | |
| Guanine nucleotide-binding protein | GNB1 | |
| N-acetylglucosamine-6-sulfatase | GNS | |
| Golgin subfamily A member 2 | GOLGA2 | |
| Golgin subfamily A member 3 | GOLGA3 | |
| Golgin subfamily A member 4 | GOLGA4 | |
| Golgin subfamily A member 5 | GOLGA5 | |
| Golgin subfamily B member 1 | GOLGB1 | |
| Golgi phosphoprotein 3 | GOLPH3 | |
| Golgi-associated PDZ and coiled-c | GOPC | |
| Golgi SNAP receptor complex mem | GOSR1 | |
| Golgi SNAP receptor complex mem | GOSR2 | |
| Glucose-6-phosphate isomerase | GPI | |
| Gelsolin | GSN | |
| Glutathione synthetase | GSS | |
| Glutathione S-transferase P | GSTP1 | |
| Histone H2B type 1-K | H2BC12 | |
| Histone H4 | H4C1 | |
| Probable histidyl-tRNA synthetase | HARS2 | |
| Histone deacetylase 6 | HDAC6 | |
| Beta-hexosaminidase subunit alph | HEXA | |
| Beta-hexosaminidase subunit beta | HEXB | |
| Protein Hook homolog 3 | HOOK3 | |
| Heat shock protein HSP 90-alpha | HSP90AA1 | |
| Heat shock protein HSP 90-beta | HSP90AB1 | |
| Endoplasmin | HSP90B1 | |
| Heat shock 70 kDa protein 4 | HSPA4 | |
| 78 kDa glucose-regulating protein | HSPA5 | |
| Heat shock cognate 71 kDa protein | HSPA8 | |

| | | |
|---|---|---|
| Heat shock protein beta-1 | HSPB1 | |
| 60 kDa heat shock protein, mitocho | HSPD1 | |
| Isoleucine-tRNA ligase, cytoplasmi | IARS | |
| Intercellular adhesion molecule 2 | ICAM2 | |
| Uncharacterised protein KIAA0947 | ICE1 | |
| Isocitrate dehydrogenase [NADP] c | IDH1 | |
| Insulin-like growth factor-binding pr | IGFBP7 | |
| Ig kappa chain C region | IGKC | |
| Acetolactate synthase-like protein | ILVBL | |
| Inosine-5'-monophosphate dehydro | IMPDH2 | |
| Insulin receptor related protein | INSRR | |
| Ras GTPase-activating-like protein | IQGAP1 | |
| Integrin alpha-5 | ITGA5 | |
| Integrin beta-1 | ITGB1 | |
| Integrin beta-5 | ITGB5 | |
| Junction plakoglobin | JUP | |
| Kinesin-like protein KIF1A | KIF1A | |
| Importin subunit beta-1 | KPNB1 | |
| Lysosome-associated membrane g | LAMP1 | |
| Lysosome-associated membrane g | LAMP1 | |
| Cytosol aminopeptidase | LAP3 | |
| Lamin-B receptor | LBR | |
| L-lactate dehydrogenase A chain | LDHA | |
| L-lactate dehydrogenase B chain | LDHB | |
| Low-density lipoprotein receptor | LDLR | |
| Vesicular integral-membrane prote | LMAN2 | |
| Leucyl-cystinyl aminopeptidase;Le | LNPEP | |
| Lipopolysaccharide-responsive and | LRBA | |
| Lysozyme g-like protein 2 | LYG2 | |
| Endoplasmic reticulum mannosyl-o | MAN1B1 | |
| Lysosomal alpha-mannosidase | MAN2B1 | |
| Epididymis-specific alpha mannosi | MAN2B2 | |
| Mitogen-activated protein kinase ki | MAP3K11 | |
| Myristoylated alanine-rich C-kinase | MARCKS | |
| S-adenosylmethionine synthase iso | MAT2A | |
| Cell surface glycoprotein MUC18 | MCAM | |
| DNA replication licensing factor MC | MCM3 | |
| Mediator of RNA polymerase II tran | MED8 | |
| Matrix Gla protein | MGP | |
| Multimerin-1 | MMRN1 | |
| Multimerin-1;Platelet glycoprotein I | MMRN1 | |
| Protein MON2 homolog | MON2 | |
| Moesin | MSN | |
| Major vault protein | MVP | |
| Myosin-9 | MYH9 | |
| Myosin-9 | MYH9 | |
| Myosin VA | MYO5A | |
| Unconventional myosin-Vc | MYO5C | |
| Myoferlin | MYOF | |
| Myoferlin | MYOF | |
| Rab effector MyRIP | MYRIP | Nightingale TD et al. 2009 |
| Rab effector MyRIP | MYRIP | |
| Rab effector MyRIP | MYRIP | |
| Nascent polypeptide associated co | NACA | |
| Alpha-N-acetylgalactosaminidase | NAGA | |
| Alpha-N-acetylglucosaminidase | NAGLU | |
| Alpha-soluble NSF attachment pro | NAPA | |
| Gamma-soluble NSF attachment p | NAPG | |
| Neurobeachin-like protein 2 | NBEAL2 | |
| NADH dehydrogenase [ubiquinone | NDUFS2 | |
| Adaptin ear-binding coat-associate | NECAP2 | |
| Neurofilament heavy polypeptide | NEFH | |
| Sialidase-1 | NEU1 | |
| Epididymal secretory protein E1 | NPC2 | |
| Neural proliferation differentiation a | NPDC1 | |
| Puromycin-sensitive aminopeptida | NPEPPSL1 | |
| Sterol-4-alpha-carboxylate 3-dehyd | NSDHL | |
| Nucleobindin-1 | NUCB1 | |
| OCIA domain-containing protein 1 | OCIAD1 | |
| Inositol polyphosphate 5-phosphata | OCRL | |
| Oxysterol-binding protein 1 | OSBP | |
| Oxysterol-binding protein-related p | OSBPL11 | |
| Ubiquitin thioesterase OTUB1 | OTUB1 | |
| Oxidation resistance protein 1 | OXR1 | |
| Protein disulfide-isomerase | P4HB | |
| Phosphofurin acidic cluster sorting | PACS1 | |
| Protein mono-ADP-ribosyltransfera | PARP4 | |
| Poly(rC)-binding protein 1 | PCBP1 | |
| Protein disulfide-isomerase A3 | PDIA3 | |
| Protein disulfide-isomerase A4 | PDIA4 | |
| Pyridoxal-dependent decarboxylas | PDXDC1 | |
| Platelet endothelial cell adhesion n | PECAM1 | |
| Platelet endothelial cell adhesion n | PECAM1 | |

| | | |
|---|---|---|
| Xaa-Pro-dipeptidase | PEPD | |
| 6-phosphofructokinase type C | PFKP | |
| 6-phosphogluconate dehydrogena | PGD | |
| Phosphoglycerate kinase 1 | PGK1 | |
| Phosphatidylinositol 4-kinase type | PI4K2A | Lopes da Silva M et al. 2016 |
| Phosphatidylinositol 4-kinase beta | PI4KB | Lopes da Silva M et al. 2016 |
| Phosphatidylinositol glycan anchor | PIGU | |
| Phosphatidylinositol-4,5-bisphosph | PIK3CA | |
| Pyruvate kinase isozymes M1/M2 | PKM | |
| Membrane-associated tyrosine- an | PKMYT1 | |
| Plakophilin-1 | PKP1 | |
| Group XV phospholipase A2 | PLA2G15 | |
| Phospholipase D1 | PLD1 | Disse J et al. 2009 |
| Phospholipase D1 | PLD1 | |
| Phospholipase D3 | PLD3 | |
| Proteolipid protein 2 | PLP2 | |
| Plasmalemma vesicle-associated | PLVAP | |
| Plexin-D1 | PLXND1 | |
| Podocalyxin | PODXL | |
| DNA-directed RNA polymerases I, | POLR2H | |
| Peptidyl-prolyl cis-trans isomerase | PPIA | |
| Protein phosphatase 1 regulatory s | PPP1R7 | |
| Protein phosphatase 2B | PPP3CB | |
| Palmitoyl-protein thioesterase 1 | PPT1 | |
| Lysosomal Pro-X carboxypeptidase | PRCP | |
| Phosphatidylinositol-3,4,5-trisphos | PREX1 | |
| DNA-dependent protein kinase cat | PRKDC | |
| Endothelial protein C receptor | PROCR | |
| Protein PRRC1 | PRRC1 | |
| 26S protease regulatory subunit 6E | PSMC4 | |
| 26S proteasome non-ATPase regu | PSMD11 | |
| Tyrosine-protein phosphatase non- | PTPN9 | |
| Peptidyl-tRNA hydrolase 2, mitoch | PTRH2 | |
| Pentraxin-related protein PTX3 | PTX3 | |
| Ras-related protein Rab-11B | RAB11B | |
| Rab11 family-interacting protein 5 | RAB11FIP5 | |
| Ras-related protein Rab-13 | RAB13 | |
| Ras-related protein Rab-14 | RAB14 | |
| Ras-related protein Rab-15 | RAB15 | |
| Ras-related protein Rab-1B;Putativ | RAB1B;RAB1C | |
| Ras-related protein Rab-27A | RAB27A | Hannah MJ et al. 2003 |
| Ras-related protein Rab-27A | RAB27A | |
| Ras-related protein Rab-32 | RAB32 | |
| Ras-related protein Rab-33B | RAB33B | |
| Ras-related protein Rab-34 | RAB34 | |
| Ras-related protein Rab-35 | RAB35 | |
| Ras-related protein Rab-37 | RAB37 | |
| Ras-related protein Rab-3A | RAB3A | Zografou S et al. 2012 |
| Ras-related protein Rab-3A | RAB3A | |
| Ras-related protein Rab-3B | RAB3B | Bierings R et al. 2012 |
| Ras-related protein Rab-3B | RAB3B | |
| Ras-related protein Rab-3D | RAB3D | Knop M et al. 2004 |
| Ras-related protein Rab-3D | RAB3D | |
| Ras-related protein Rab-3D | RAB3D | |
| Ras-related protein Rab-6A | RAB6A | |
| Ras-related protein Rab-7a | RAB7A | |
| Rac family small GTPase 1 | RAC1 | |
| Guanine nucleotide-binding protein | RACK1 | |
| Ras-related protein Ral-A | RALA | |
| RAS like proto-oncogene A | RALA | |
| Ral guanine nucleotide dissociation | RALGDS | |
| Ras-related protein 1 | RAP1B | |
| EPAC/Rap guanine nucleotide exc | RAPGEF3 | |
| Ras-interacting protein | RASIP1 | |
| Protein RER1 | RER1 | |
| Rhomboid domain-containing prote | RHBDD2 | |
| Transforming protein RhoA | RHOA | |
| Ras homolog family member A | RHOA | |
| Rho-related GTP-binding protein R | RHOC | |
| RAB6A-GEF complex partner prote | RIC1 | |
| Regulator of microtubule dynamics | RMDN3 | |
| Ribonuclease inhibitor | RNH1 | |
| Roundabout homolog 4 | ROBO4 | |
| 60S ribosomal protein L7 | RPL7 | |
| 40S ribosomal protein S20 | RPS20 | |
| 40S ribosomal protein SA | RPSA | |
| S100 Calcium-Binding Protein A10 | S100A10 | |
| Phosphatidylinositide phosphatase | SACM1L | |
| Secretory carrier-associated memb | SCAMP1 | |
| Lysosome membrane protein 2 | SCARB2 | |
| Sec1 family domain-containing pro | SCFD1 | |
| Secernin-1 | SCRN1 | |
| Vesicle-trafficking protein SEC22b | SEC22B | |

| | | |
|---|---|---|
| Protein transport protein Sec23A | SEC23A | |
| Protein transport protein Sec23B | SEC23B | |
| SEC23-interacting protein | SEC23IP | |
| Protein transport protein Sec24B | SEC24B | |
| Protein transport protein Sec24C | SEC24C | |
| P-selectin | SELP | Bonfanti R et al. 1989 |
| P-selectin | SELP | |
| Serpin B12 | SERPINB12 | |
| Serpin B6 | SERPINB6 | |
| Serpin B9 | SERPINB9 | |
| Plasminogen activator inhibitor 1 | SERPINE1 | |
| Serpin H1 | SERPINH1 | |
| Protein SET | SET | |
| 14-3-3 protein sigma | SFN | |
| Vesicle transport protein SFT2C | SFT2D3 | |
| N-sulphoglucosamine sulphohydro | SGSH | |
| Zinc transporter 5 | SLC30A5 | |
| Zinc transporter 6 | SLC30A6 | |
| Zinc transporter 7 | SLC30A7 | |
| CMP-sialic acid transporter | SLC35A1 | |
| Solute carrier family 35 member E1 | SLC35E1 | |
| Mothers against decapentaplegic h | SMAD1 | |
| Synaptosome associated protein 2 | SNAP23 | |
| Synaptosomal-associated protein 2 | SNAP29 | |
| U5 small nuclear ribonucleoprotein | SNRNP200 | |
| SPARC | SPARC | |
| Protein spire homolog 1 | SPIRE1 | |
| Lupus La protein | SSB | |
| Erythrocyte band 7 integral membr | STOM | |
| Syntaxin-12 | STX12 | |
| Syntaxin-16 | STX16- NPEPL1;STX16 | |
| Syntaxin-3 | STX3 | Schillemans M et al. 2018 |
| Syntaxin-4 | STX4 | |
| Syntaxin-5 | STX5 | |
| Syntaxin-7 | STX7 | |
| Syntaxin-binding protein 1 | STXBP1 | van Breevoort D et al. 2014 |
| Syntaxin-binding protein 1 | STXBP1 | |
| Munc18-c | STXBP3 | |
| Syntaxin-binding protein 5 | STXBP5 | |
| Sin3 histone deacetylase corepres | SUDS3 | |
| SUN domain-containing protein 2 | SUN2 | |
| Synaptotagmin-like protein 4 | SYTL4 | Bierings R et al. 2012 |
| Synaptotagmin-like protein 4-a | SYTL4 | |
| Transgelin-2 | TAGLN2 | |
| Transaldolase | TALDO1 | |
| TBC1 domain family member 22A | TBC1D22A | |
| Transferrin receptor protein 1;Tran | TFRC | |
| Protein-glutamine gamma-glutamy | TGM2 | |
| Protein-glutamine gamma-glutamy | TGM3 | |
| Trans-Golgi network integral memb | TGOLN2 | |
| Thrombospondin-1 | THBS1 | |
| Thrombospondin-1 | THBS1 | |
| Tight junction-associated protein 1 | TJAP1 | |
| Transketolase | TKT | |
| Transmembrane 9 superfamily me | TM9SF3 | |
| Transmembrane emp24 domain-co | TMED5 | |
| Transmembrane protein 2 | TMEM2 | |
| Transmembrane protein 43 | TMEM43 | |
| TATA element modulatory factor | TMF1 | |
| Thioredoxin-related transmembran | TMX1 | |
| Toll-interacting protein | TOLLIP | |
| Mitochondrial import receptor subu | TOMM70A | |
| Torsin-1A-interacting protein 1 | TOR1AIP1 | |
| Triosephosphate isomerase | TPI1 | |
| Tropomyosin alpha-4 chain | TPM4 | |
| Tripeptidyl-peptidase 1 | TPP1 | |
| Thyroid receptor-interacting protein | TRIP11 | |
| Tubulin alpha-1B chain | TUBA1B | |
| Tubulin alpha-1C chain | TUBA1C | |
| Tubulin beta chain | TUBB | |
| Tubulin beta-2A chain | TUBB2A | |
| Tubulin beta-3 chain | TUBB3 | |
| Tubulin beta-2C chain | TUBB4B | |
| Tubulin beta-6 chain | TUBB6 | |
| Thioredoxin domain-containing pro | TXNDC5 | |
| Thioredoxin reductase 1, cytoplasm | TXNRD1 | |
| UDP-N-acetylhexosamine pyropho | UAP1L1 | |
| Ubiquitin -like modifier-activating e | UBA1 | |
| Ubiquitin-associated protein 2-like | UBAP2L | |
| Polyubiquitin-C | UBC | |
| Ubiquitin-conjugating enzyme E2 J | UBE2J1 | |
| Ceramide glucosyltransferase | UGCG | |
| UDP-glucose 6-dehydrogenase | UGDH | |

| | | |
|---|---|---|
| Protein unc-13 homolog B/Munc13 | UNC13B | |
| Munc13-4 | UNC13D | |
| General vesicular transport factor p | USO1 | |
| Vesicle associated membrane prot | VAMP3 | |
| Vesicle-associated membrane prot | VAMP3;VAMP2 | Pulido IR et al. 2011 |
| Vesicle associated membrane prot | VAMP8 | |
| Synaptic vesicle membrane protein | VAT1 | |
| Vinculin | VCL | |
| Transitional endoplasmic reticulum | VCP | |
| Vimentin | VIM | |
| Spermatogenesis-defective protein | VIPAS39 | |
| Vacuolar protein sorting-associated | VPS13B | |
| Vacuolar protein sorting-associated | VPS45 | |
| Vacuolar protein sorting-associated | VPS51 | |
| V-set and immunoglobulin domain- | VSIG8 | |
| Vesicle transport through interactio | VTI1A | |
| von Willebrand factor A domain-co | VWA5B1 | |
| von Willebrand factor | VWF | |
| Tryptophanyl-tRNA synthesase, cy | WARS1 | |
| WD repeat-containing protein 1 | WDR1 | |
| WD repeat-containing protein 11 | WDR11 | |
| Exportin-1 | XPO1 | |
| Nuclease-sensitive element bindin | YBX1 | |
| Protein YIF1A | YIF1A | |
| Protein YIF1B | YIF1B | |
| Protein YIPF4 | YIPF4 | |
| Protein YIPF5 | YIPF5 | |
| 14-3-3 protein beta/alpha | YWHAB | |
| 14-3-3 protein epsilon | YWHAE | |
| 14-3-3 protein zelta/delta | YWHAZ | |
| Palmitoyltransferase ZDHHC13 | ZDHHC13 | |
| Palmitoyltransferase ZDHHC17 | ZDHHC17 | |
| Zinc finger protein-like 1 | ZFPL1 | |
| Zinc finger protein 598 | ZNF598 | |
| Heat shock 70 kDa protein 1A/1B | | |
| Ig lambda-2 chain C regions | | |
